# Unraveling a Bacterial Starvation Response Through the Direct Targets of a Starvation-Induced Transcriptional Activator

**DOI:** 10.1101/2021.11.28.470282

**Authors:** Muqing Ma, Ting Li, David J. Lemon, Eduardo A. Caro, Linnea Ritchie, Charles Ryan, Victoria M. Spearing, Roy D. Welch, Anthony G. Garza, Kimberly A. Murphy

## Abstract

Organisms frequently encounter environments with nutrient shortages and their survival depends on changes in physiology and the ability to conserve resources. In bacteria, many physiological changes associated with starvation have been identified, but the underlying genetic components and regulatory networks that direct these physiological changes are often poorly defined. Here, we aimed to better define the gene regulatory networks that mediate the starvation response in *Myxococcus xanthus*, a bacterium that copes with starvation by producing fruiting bodies filled with dormant and stress-resistant spores. We focused on the direct promoter/gene targets of Nla28, a transcriptional activator/enhancer binding protein (EBP) that is important for early rounds of gene expression following starvation. Using expression profiling to identify genes that are downregulated in *nla28* mutant cells and bioinformatics to identify the putative promoters of these genes, 12 potential promoter targets (37 genes) of Nla28 were identified. The results of in vitro promoter binding assays, coupled with in vitro and in vivo mutational analyses, suggested that the 12 promoters are in vivo targets of Nla28 and that Nla28 dimers use tandem, imperfect repeats of an 8-bp sequence for binding. Interestingly, nine of the Nla28 target promoters are intragenic, located in the protein coding sequence of an upstream gene or in the protein coding sequence of one gene within an operon (internal promoters). Based on mutational analyses, we concluded that the 12 Nla28 target loci contain at least one gene important for production of stress-resistant spores following starvation. Most of these loci contain genes predicted to be involved in regulatory or defense-related functions. Using the consensus Nla28 binding sequence, followed by bioinformatics and expression profiling, 58 additional promoters and 102 genes were tagged as potential Nla28 targets. Among these putative Nla28 targets, functions such as regulatory, metabolic and cell envelope biogenesis were commonly assigned to genes.

## INTRODUCTION

Bacteria typically have numerous transcription factors to activate and/or repress transcription of genes that are important for development. In the soil bacterium *Myxococcus xanthus*, the early developmental pathway relies heavily on enhancer binding proteins or EBPs (1), which are transcriptional activators that help σ^54^ -RNA polymerase initiate transcription at σ^54^-type promoters (2–4). Indeed, a cascade of EBPs modulates changes in developmental gene transcription during sequential stages of early development (5). The aim of this study was to identify and characterize the developmental gene targets of an early-functioning EBP called Nla28.

During development, *M*. *xanthus* forms a biofilm that contains a mat of rod-shaped cells, known as peripheral rods (6, 7), and multicellular fruiting bodies that house about 100,000 dormant and stress-resistant spores (8). Starvation triggers the developmental process, and much is known about the subsequent morphological changes that yield spore-filled fruiting bodies. Namely, cells migrate into an aggregation center, the cell aggregate becomes larger and eventually develops the dome-shaped appearance of a fruiting body, the rod-shaped cells in the newly formed fruiting body differentiate into spherical cells, and the spherical cells mature into stress-resistant spores.

In standard developmental assays, at least two starvation-induced signaling events must occur before cells begin building fruiting bodies. The first event is the RelA-mediated accumulation of the intracellular starvation signal (p)ppGpp (9, 10). The second event, which depends on relatively high levels of (p)ppGpp (10), is the accumulation of an extracellular signal called A-signal. A-signal is a cell-density signal composed of amino acids and perhaps peptides (11–13). Thus, (p)ppGpp accumulation indicates that cells are starving, and accumulation of A-signal indicates that enough starving cells are present to build a fruiting body.

Our early work linked four of the 53 *M*. *xanthus* EBPs to the accumulation of early developmental signals (14–18). The EBP Nla28, which is the focus of this study, was linked to A-signal production via extracellular complementation assays (14). Nla28 is a response regulator-type EBP that partners with the membrane-bound histidine protein kinase Nla28S to form a two component signal transduction system (19, 20). Two pieces of data led to the suggestion that the Nla28S/Nla28 signal transduction system might respond to A-signal, in addition to being important for A-signal production (20). First, *nla28S* and *nla28* form a two-gene operon and Nla28 is important for autoregulation (5). Secondly, A-signal is important for full developmental expression of the *nla28S* gene and presumably *nla28S*-*nla28* operon (19).

In addition to being connected to A-signal production, *nla28* is known to be expressed early in development and important for expression of early developmental genes; inactivation of *nla28* impairs or abolishes expression of many genes that are induced 1-2 hours post-starvation (5). This finding led to the suggestion that the Nla28/Nla28 signal transduction system targets starvation-associated or stress-responsive genes (18, 20). Indeed, the results of a recent study suggest that Nla28 directly modulates the activities of three natural product promoters during the transition into stationary phase and during development; both events are associated with nutrient depletion (21).

Although Nla28 is important for expression of hundreds of *M*. *xanthus* developmental genes and for the developmental process (5, 14), only two classes of developmental genes have been linked to direct Nla28-mediated regulation: regulatory genes and natural product-associated genes (5, 21). One goal of this study was to better understand how Nla28 identifies its target promoters and to better understand how these promoters are arranged. Another goal was to identify additional developmental promoter targets of Nla28, which would in turn allow us to analyze the developmental genes under direct Nla28 control and hence to better understand the function of Nla28.

Since EBPs modulate transcription at σ^54^ promoters (22, 23), we focused our search for direct Nla28 targets on known or putative σ^54^ promoters located upstream of developmentally regulated single genes and operons (5). In the initial experiments, we confirmed twelve potential σ^54^ promoter targets of Nla28 and identified the likely promoter binding sites of Nla28 using expression data, in vitro promoter binding assays, and in vitro and in vivo mutational analyses. As in previous studies of Nla EBPs (5, 18, 21), our analysis placed the vast majority of the Nla28 target promoters inside genes and not in intergenic regions. Several of the previously uncharacterized targets of Nla28 were subsequently analyzed via insertion mutagenesis and all were linked to production of stress-resistant spores during starvation-induced development. Among the confirmed Nla28 target loci, genes predicted to be involved in regulatory or defense-related functions were common. An additional 58 putative σ^54^ promoters and 102 genes were tagged as potential targets of Nla28 using the consensus Nla28 binding sequence [CT(C/G)CG(C/G)AG]. Among these putative Nla28 targets, functions such as regulatory, metabolic and cell envelope biogenesis were commonly assigned to genes.

## MATERIALS AND METHODS

### Bacterial strains and plasmids

The strains, plasmids and primers used in this study are shown in Tables S1 and S2. Plasmids generated for this study were analyzed via DNA sequence analysis. *M. xanthus* strains were confirmed as described below. Plasmid insertions in Nla28 target genes in wild-type strain DK1622 were generated as previously described (14, 24) and confirmed via PCR.

### Growth and development

*Escherichia coli* strains were grown at 37^°^C in LB broth containing 1.0% tryptone, 0.5% yeast extract, and 0.5% NaCl or on plates containing LB broth and 1.5% agar. LB broth and LB plates were supplemented with 40 μg of kanamycin sulfate/ml, 100 μg of ampicillin/ml or 10 μg of oxytetracycline/ml as needed.

*M. xanthus* strains were grown at 32^°^C in CTTYE broth [1.0% Casitone, 0.5% yeast extract, 10 mM Tris-HCl (pH 8.0), 1 mM KH_2_PO_4_, and 8 mM MgSO_4_] or on plates containing CTTYE broth and 1.5% agar. CTTYE broth and plates were supplemented with 40 μg of kanamycin sulfate/ml or 10 μg of oxytetracycline/ml as needed.

Fruiting body development was induced by placing *M. xanthus* cells on plates containing TPM buffer [10 mM Tris-HCl (pH 8.0), 1 KH_2_PO_4_, and 8 mM MgSO_4_] and 1.5% agar or in 6-well microtiter plates containing MC7 buffer (10 mM MOPS, 1 mM CaCl_2_, final pH 7.0), and incubating the plates at 32°C. Briefly, *M*. *xanthus* strains were grown in flasks containing CTTYE broth and the cultures were incubated at 32°C with vigorous swirling. The cells were pelleted when the cultures reached a density of about 5 × 10^8^ cells/ml, the supernatants were removed, and the cells were resuspended in TPM buffer or MC7 buffer to a final density of 5 × 10^9^ cells/ml. Aliquots of the cells in TPM buffer were spotted onto TPM agar plates and aliquots of the cells in MC7 buffer were placed in 6-well microtiter plates containing MC7 buffer. The cells were incubated at 32°C and development was monitored as previously described (14, 18). Cells were harvested at various times during development and prepared for quantitative PCR (qPCR), β-Galactosidase assays or sporulation assays.

To examine the sporulation efficiency of each *M*. *xanthus* strain, developing cells were harvested from TPM agar plates after 5 days and the cells were placed in 400 μl of TPM buffer. Aliquots of the cell suspension were placed on a Petroff-Hausser counting chamber and phase-contrast microscopy was used to determine the number of spherical-shaped cells that were present. Other aliquots of the cell suspension were sonicated, and the sonicated cells were incubated at 50°C for 2 hours. The number of heat- and sonication-resistant spores that germinated into colonies was determined by placing heat- and sonication-treated cells in liquified CTT soft agar [1.0% Casitone, 10 mM Tris-HCl (pH 8.0), 1 mM KH_2_PO_4_, 8 mM MgSO_4_ and 0.7% agar], pouring the soft agar onto CTTYE agar plates and incubating the plates at 32°C for 5 days. Similar assays were performed on spores formed when glycerol was added to CTTYE broth cultures of *M*. xanthus cells (final glycerol concentration of 0.5 M).

### Motility assays

Motility assays were performed as previously described (14, 24). Briefly, *M*. *xanthus* cells were grown to a density of about 5 × 10^8^ cells/ml in CTTYE broth. The cells were pelleted by centrifugation, the supernatant was removed, and the cells were resuspended in CTTYE broth to a density of 5 × 10^9^ cells/ml. Aliquots (3 μl) of the cell suspensions were spotted onto CTTYE plates containing 1.5 or 0.4% agar, the spots were allowed to dry, and the plates were placed at 32°C. After 3 days of incubation, five swarms of each strain were measured, and their mean diameter was normalized to the mean diameter of five swarms formed by wild-type strain DK1622.

### Plasmid transfer to *M*. *xanthus*

Plasmids containing internal fragments of Nla28 target genes or wild-type or mutant copies of the *nla28* promoter were electroporated into wild-type strain DK1622 as described previously (25). Kan^r^ electroporants that contained a plasmid integrated in the *nla28* locus or in a *nla28* target gene by homologous recombination, or containing a plasmid integrated in the chromosomal Mx8 phage attachment site (*attB*) by site-specific recombination were identified via PCR or PCR and DNA sequencing. Kan^r^ electroporants carrying a single plasmid insertion were assayed for development, motility or promoter activity as needed.

### Standard DNA procedures

Chromosomal DNA from wild-type *M. xanthus* strain DK1622 was extracted using a ZYMO Research gDNA extraction kit. Oligonucleotides used in PCR reactions were synthesized by Integrated DNA Technologies (IDT) and are listed in Table S2. Plasmid DNA was extracted using the Promega Nucleic Acid Purification kit. Amplified and digested DNA fragments were purified using the Gel Extraction Minipreps kit from Bio Basic. For all kits, the manufacturer’s protocols were used. The compositions of all plasmids and promoter fragments were confirmed by DNA sequencing (Genewiz).

### In vivo mutational analysis of the putative Nla28 binding site in the *nla28* promoter

A purified, 494-bp DNA fragment of the *nla28* promoter region containing the putative σ^54^-RNA polymerase binding site and the Nla28 binding site was cloned into the pCR-Blunt vector (Invitrogen). Site-directed mutations in the putative Nla28 binding site were generated using the Quickchange system (Agilent) as previously described (5, 18), and the promoter mutations were confirmed by DNA sequence analysis. The wild-type and mutant promoter fragments were subsequently cloned into promoterless *lacZ* expression plasmids, which were designed to generate transcriptional fusions between the cloned promoters and the *lacZ* gene. One of these plasmids, pREG1727, confers kanamycin resistance (Kan^r^) and carries the Mx*8 attP* site, allowing for site-specific integration at the Mx*8* phage attachment site *attB* in the chromosome of *M*. *xanthus* (26). The other plasmid (pTL1) is a derivative of pREG1727 with *attP* removed; this plasmid is unable to integrate at *attB* but is capable of recombining at the native promoter locus via the homologous cloned promoter fragment.

The *nla28* promoter-*lacZ* fusion plasmids were introduced into wild-type strain DK1622, and Kan^r^ isolates carrying a plasmid integrated at the Mx*8* phage attachment site in the chromosome or at the native *nla28* locus in the chromosome were identified via PCR and analyzed via DNA sequencing. Cells were prepared for development on TPM agar as described above, then allowed to develop for various amounts of time before being harvested and quick-frozen in liquid nitrogen. β-Galactosidase assays, which were used to infer promoter activities, were performed on quick-frozen cell extracts as previously described (21, 27). β-Galactosidase-specific activity is defined as nanomoles of o-nitrophenol (ONP) produced per minute per milligram of protein. β Galactosidase-specific activity of the wild-type and mutant promoters at each time point was analyzed in triplicate.

### Expression and purification of Nla28-DBD

A fragment of the *nla28* gene corresponding to the Nla28 DNA Binding Domain (Nla28-DBD) (5, 21) was PCR amplified using gene-specific primers (Table S2) and then cloned into plasmid pMAL-c5x to create an N-terminal maltose binding protein (MBP) fusion to Nla28-DBD. The Nla28-DBD expression plasmid was introduced into *E*. *coli* strain BL21 (DE3) using electroporation. Cells containing Nla28-DBD expression plasmids were grown in rich LB broth to a density of approximately 2 × 10^8^ cells/ml. Protein expression was induced by the addition of 0.3 mM Isopropyl β-D-1 thiogalactopyranoside (IPTG) to the culture and the subsequent incubation of the culture for 12 hours at 15°C. Cells were pelleted via centrifugation and resuspended in 25 ml column buffer (20 mM Tris-HCl, 200 mM NaCl, 1 mM EDTA, 5 U/ml DNase I, 1 mM DTT, pH 7.4) per liter of culture. The resuspended cells were lysed by a combination of freeze-thawing and sonication, and pelleted by centrifugation. The crude extract (supernatant) containing Nla28-DBD was diluted by adding 125 ml of cold column buffer to every 25 ml aliquot of crude extract. For purification of Nla28-DBD from the diluted crude extract, diluted crude extract containing Nla28-DBD was loaded onto 5 ml MBPTrap HP columns (GE Healthcare) at a flow rate of 5 ml/min and washed with 600 ml cold column buffer at a flow rate of 10 ml/min on an ÄKTA Fast Protein Liquid Chromatography (FPLC) system (GE Healthcare). Nla28-DBD was eluted using 100 ml cold column buffer containing 10 mM maltose; the flow rate was 5 ml/min and 20 fractions containing 5 ml were collected. The presence of eluted Nla28-DBD was detected by UV absorbance at 280 nm. Nla28-DBD-containing fractions were pooled and incubated with 1 mg of Factor Xa at 4°C overnight to cleave the MBP tag. Nla28-DBD was separated from MBP and concentrated to about 1 mg/ml using Amicon Ultra centrifugal filter units (EMD Millipore). SDS-PAGE and Bradford assays were used to determinate the purity and concentration of Nla28-DBD.

### Electrophoretic mobility shift assays (EMSAs)

The PCR-generated fragments of the putative Nla28 target promoters contained approximately 140-185 bp of DNA upstream of the σ^54^-RNA polymerase binding sites, which were identified experimentally (5, 18, 28, 29) or using a bioinformatics tool (PromScan) that was specifically developed to find such sites in the sequences of bacterial DNA (30). For use in electrophoretic mobility shift assays, the promoter fragments were PCR amplified using 5’Cy5-labeled primers synthesized by IDT (Table S2). Binding reactions contained EMSA buffer (25□mM Tris/acetate [pH 8.0], 8.0□mM magnesium acetate, 10□mM KCl, 1.0□mM DTT), 2.0 μM, 1.5 μM, 1.0 μM, 0.75 μM, 0.5 μM, or 0.25 μM of Nla28-DBD and 1 ng of a 5’ Cy5-labeled promoter fragment. The binding reactions were allowed to proceed for 30 minutes at 30°C, and the binding reactions were analyzed using native (non-denaturing) PAGE and a Bio-Rad imager.

The affinity of Nla28-DBD binding to target promoters was assessed by performing EMSAs with various concentrations of Nla28-DBD. For each promoter fragment, the titration series was performed in triplicate and the signal intensities of shifted and unshifted complexes were determined using Fiji ImageJ (31). The percentage of shifted complex (Nla28-DBD-bound DNA) for each concentration of Nla28-DBD in each replicate was calculated as follows: signal intensity of shifted complex / intensities of shifted complex + unshifted free DNA.

### Quantitative PCR (qPCR)

To examine expression of Nla28 target genes, wild-type cells and *nla28* mutant cells were harvested during vegetative growth (0 hours) and during development in submerged cultures. Total cellular RNA was isolated from developmental cells using the RNAprotect Bacteria Reagent (Qiagen) and the RNeasy Mini Kit (Qiagen) as described in the manufacturer’s protocols. To help lyse developmental cells, 0.1 mm diameter glass beads were added after the lysis buffer and the cell suspensions were subjected to vigorous shaking using a VWR DVX-2500 multi-tube vortexer. For each time point (0, 1, 2, 8, 12 or 24 hours) total RNA was isolated from seven independent wild-type samples and from seven independent *nla28* mutant samples. The wild-type RNA samples from each time point were pooled and the *nla28* mutant RNA samples from each time point were pooled, and the pooled samples were subsequently used to generate cDNA as previously described (18, 32). The CFX Connect real-time PCR detection system (Bio-Rad) was used to perform the qPCR analysis (18). Each time point for the wild-type and *nla28* mutant was analyzed in triplicate. Relative fold-changes in mRNA levels were calculated using the reference gene *rpoD* or 16S rRNA and the ΔΔCT method as previously described (18, 32).

### Identifying potential targets of Nla28

To identify direct targets of Nla28, we started with a set of *M*. *xanthus* genes that were previously classified as developmentally regulated; expression of the genes increased at least 2-fold during development [our DNA microarray data on Gene Expression Omnibus, accession # GSE13523] (5). We then searched for developmentally regulated genes that had at least a 2-fold decrease in expression in a *nla28* mutant relative to that in wild-type cells (5). Fifty-one developmentally regulated genes met the criterion and hence were classified as Nla28-dependent developmental genes. We should note that the genes in the *actB*, *nla6* and *nla28* operons were not included in this analysis, as the results of previous work indicated that the certain regions of their promoters are bound by Nla28-DBD; thus, these operons were already known to be good candidates for direct in vivo regulation by Nla28 (5).

The DNA region upstream of the 51 Nla28-dependent genes were scanned for putative σ^54^ promoters using the algorithm developed for PromScan, a bioinformatics tool that identifies potential σ -RNA polymerase binding sites using conserved nucleotides in the -12 bp and -24 bp regions of σ^54^ promoters (30). Because of the distant location (in the primary DNA structure) of the σ^54^ promoter elements of the *nla6* and *nla28* operons, we searched for putative σ^54^ promoters in the 500-bp region upstream of each Nla28-dependent developmental gene. Intergenic and intragenic regions were evaluated, since previous mutational analyses placed the putative σ^54^-RNA polymerase binding sites in the *actB* and *nla6* promoters in intragenic regions and the putative σ^54^- RNA polymerase binding site in the *nla28* promoter in an intergenic region (Figure S1). We should note that in previous trial runs with known σ^54^ promoters and non-σ^54^ promoters, PromScan yielded a false positive rate of about 4% and a false negative rate of about 19% (5). Therefore, we anticipated similar false assignments in our search.

Of 51 Nla28-dependent developmental genes that were evaluated as described above, only nine had a putative σ^54^-RNA polymerase binding site in the 500-bp region. We should note that one of the putative σ^54^-RNA polymerase binding sites was previously identified during the characterization of the *pilA* promoter region (28). These nine promoters, along with the promoters of the *actB*, *nla6* and *nla28* operons (see Table 1), were considered candidates for direct in vivo regulation by Nla28. Electrophoretic mobility shift assays were subsequently used to examine whether Nla28-DBD was capable of binding to at least one DNA fragment in each promoter region; the fragments correspond to DNA flanking the putative σ^54^-RNA polymerase binding sites in the -12 and -24 bp- regions. Nla28-DBD was consistently positive for binding to at least one fragment of each promoter region and, as shown in Table 1, similar 8-bp direct repeat sequences were found in all these fragments. The consensus half binding site for Nla28 dimers derived from these repeat sequences is CT(C/G)CG(C/G)AG. Since EBP dimers bind to tandem repeats typically located well upstream of the -12 and -24 bp-regions of σ^54^-type promoters (22, 23), we examined whether the 300-bp of DNA located upstream of the putative -12 and -24-bp promoter regions contained additional sequences closely matching the consensus half binding site of Nla28. As shown in Figure S1, additional (putative) half binding sites for Nla28 were found in many of the promoter regions.

**TABLE 1.**
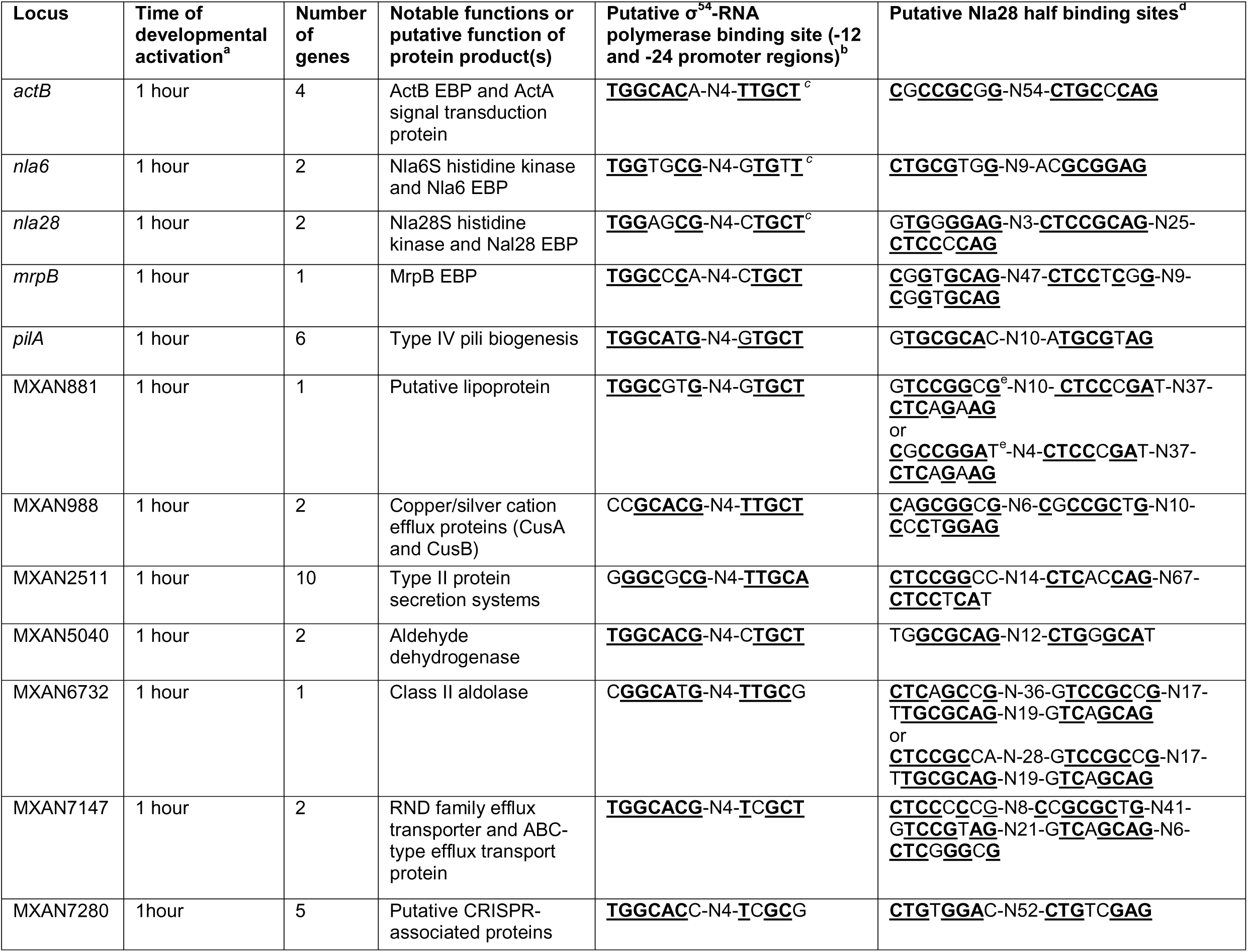
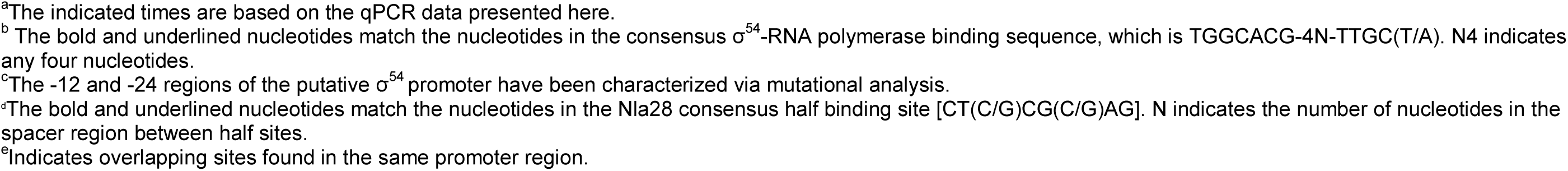
Potential developmental targets of the EBP Nla28

In a subsequent search for additional Nla28 targets, the *M*. *xanthus* genome sequence (15) was scanned for close matches to the consensus Nla28 half binding site. In particular, we looked for 8-bp sequences with no more than two mismatches relative to the consensus Nla28 half binding site. Since EBP dimers bind to DNA, we also looked for potential Nla28 half sites arranged in tandem; tandem sites were defined as putative Nla28 half sites separated by no more than 25 bp, which was the mean number of base pairs between putative Nla28 half sites in the 12 promoter regions we characterized. We then searched for putative σ^54^-RNA polymerase binding sites downstream of the tandem sequences. The locations and sequences of putative Nla28 half binding sites and σ^54^-RNA polymerase binding sites are shown in Table S3. Since we were interested in Nla28’s developmental gene targets and also developmental genes whose expression shows a strong dependence on Nla28, we further curated the list of potential Nla28 targets by doing the following: 1) we identified single genes and operons located downstream of the putative σ^54^-RNA polymerase binding sites in the -12 and -24-bp regions; 2) using DNA microarray data (5), we determined which of the single genes and operons showed at least a 2-fold increase in expression during the development of wild-type cells; 3) using DNA microarray data (5), we also identified the single genes and operons that showed at least a 2-fold decrease in expression during the development of *nla28* mutant cells. The single genes and operons that met the above criteria are shown in Table S4.

### Statistical analysis

Individual sample sizes are specified in each figure legend. Comparisons between groups were assessed by two-way analysis of variance (ANOVA) with Tukey’s multiple comparisons *post hoc* tests, as appropriate. The significance level was set at p<0.05 or lower. Prism (GraphPad) v9.2 was used for all analyses.

## RESULTS

### Identifying a set of potential developmental promoter targets of Nla28

In our initial studies, we aimed to identify and characterize a collection of developmental promoters that are likely to be directly regulated by Nla28 in vivo. As described in the *Materials and Methods*, nine developmental promoters were classified as potential targets of Nla28 using DNA microarray expression data (5) and bioinformatics (30). Three additional developmental promoters, namely the promoters of the EBP gene operons *actB*, *nla6* and *nla28*, were added to the collection of nine putative Nla28 targets because of previous data indicating that they are likely Nla28 targets (5, 18). As noted above, the putative σ^54^-RNA polymerase binding regions of the *actB*, *nla6* and *nla28* promoters were characterized via in vivo mutational analyses (5, 29) and the putative σ -RNA polymerase binding regions of the *pilA* promoter were identified via primer extension analysis (28). In contrast, the putative σ^54^-RNA polymerase binding sites and promoter regions of the nine newly identified targets of Nla28 have not been characterized.

The locations and sequences of the putative σ^54^-RNA polymerase binding sites in the -12- and -24-bp regions of the 12 promoters are shown in Figure S1 and Table 1, respectively. It is noteworthy that the 9/12 of the putative σ^54^-RNA polymerase binding sites or core σ^54^ promoter regions are located within genes and not in intergenic sequences. This finding is consistent with previous studies that placed many core σ^54^ promoter regions in the coding sequences of *M*. *xanthus* genes (5, 15, 18, 21) and raises the possibility that intragenic σ^54^ promoters might be common in *M*. *xanthus* and in bacteria in general. As in previous analyses of *M*. *xanthus* σ promoters, some of the intragenic -12- and -24-bp regions are located in the protein coding sequence of an upstream gene (upstream promoters) and some are located in the protein coding sequence of one gene in an operon (internal promoters). The implications of these findings and the findings of other studies are addressed in the *Discussion*.

### Promoter fragments that are positive for in vitro Nla28-DBD binding have similar 8-bp repeats

In preliminary electrophoretic mobility shift assays (EMSAs), we examined the ability of purified Nla28-DBD (Nla28 DNA binding domain) to bind to DNA fragments flanking the putative -12- and -24-bp regions of the 12 promoters. At least one fragment of each promoter region was consistently positive for Nla28-DBD binding. The results of EMSAs performed with 2 μM of Nla28-DBD and 5’ Cy5-labeled versions of the promoter fragments are shown in Figure 1. All the promoter fragments except the negative control *dev* promoter fragment, which is a fragment of a non-σ^54^ promoter, produced at least one shifted complex; however, a number of the promoter fragments produced more than one shifted complex, as discussed below.

**Fig 1.**
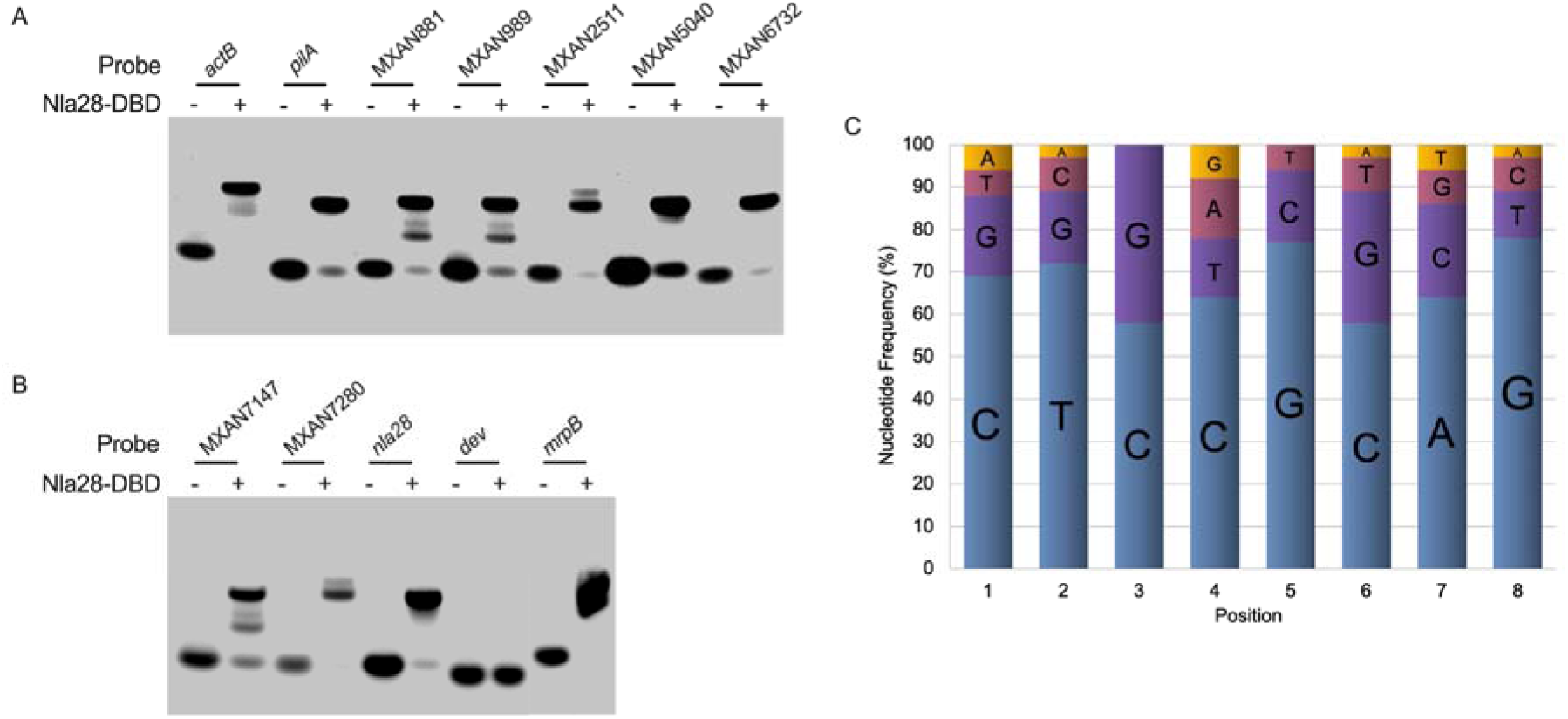
Promoter fragments that are positive for in vitro Nla28-DBD binding have similar 8-bp repeats. (A and B) Purified Nla28-DBD binds to fragments of 11 developmental promoters. EMSAs performed with purified Nla28-DBD and an *actB*, *pilA*, MXAN881, MXAN989, MXAN2511, MXAN5040, MXAN6732 (A), MXAN7147, MXAN7280, *nla28*, *dev* or *mrpB* (B) promoter fragment containing putative Nla28 binding sites. Binding reactions were performed with (+) or without (−) 2 μM of purified Nla28-DBD and a Cy5 5’ end-labeled promoter fragment flanking the -12- and-24-bp regions. (C) Nucleotide frequency at each position of the putative Nla28 half binding sites. Similar 8-bp sequences were found in all promoter fragments that were positive for Nla28 binding. The 8-bp sequences were used to generate nucleotide frequency data and to generate the following consensus 8-bp sequence or consensus Nla28 half binding site: CT(C/G)CG(C/G)AG.

When we examined the DNA sequences in the fragments that were positive for Nla28-DBD binding, we identified at least two, but often 3-5 8-bp sequences that are similar (Table 1). Since EBP dimers typically bind to tandem repeat sequences or tandem half binding sites, we concluded that at least two of the 8-bp sequences in each promoter fragment are likely to serve as Nla28-DBD half binding sites. We also speculated that Nla28-DBD might be able to form different binding complexes when the promoter fragments have more than one half site pair; this might explain why promoter fragments such as MXAN881, MXAN988 and MXAN7147 yielded more than one shifted complex in EMSAs (Figure 1). Of course, some of the 8-bp sequences may not be Nla28-DBD half binding sites or may be low affinity half binding sites, and some of the pairs of half binding sites might be too distant (in the primary sequence) for efficient binding of an Nla28-DBD dimer. Hence, at this point it is difficult predict the number and types of complexes that Nla28-DBD will be able to form with each promoter fragment. We should note that many of the promoter regions contain a second putative cluster of Nla28 half sites (Figure S1); we identified these sites using the consensus Nla28 half binding site 5’-CT(C/G)CG(C/G)AG-3’ (see Figure 1C).

### Two 8-bp sequences in the *actB* promoter and two in the *nla28* promoter are crucial for in vitro Nla28-DBD binding

Since all the promoter fragments that were positive for Nla28-DBD binding in EMSAs contained similar 8-bp sequences, we asked if these sequences are important for Nla28-DBD binding. We initially focused on the *actB* promoter fragment, which has only two 8-bp sequences or two putative half sites for binding of a Nla28-DBD dimer. Three *actB* promoter fragments were generated for the in vitro Nla28-DBD binding analysis (see Figure 2A). One of the *actB* promoter fragments had wild-type half sites 1 and 2, one fragment had a wild-type half site 1 and a half site 2 that was converted to all A nucleotides, and one fragment had wild-type half site 2 and a half site 1 that was converted to all A nucleotides. The results of the EMSAs performed with 2 μM of Nla28-DBD and 5’ Cy5-labeled versions of wild-type and mutant promoter fragments are shown in Figure 2B. As in previous EMSAs, Nla28-DBD was capable of binding to the *actB* promoter fragment carrying two wild-type half sites. However, no binding was detected when either half site 1 or half site 2 contained all A nucleotides. This finding is consistent with our proposal that the two 8-bp sequences serve as half binding sites for Nla28-DBD dimers and that both sites are important for Nla28-DBD binding.

**Fig 2.**
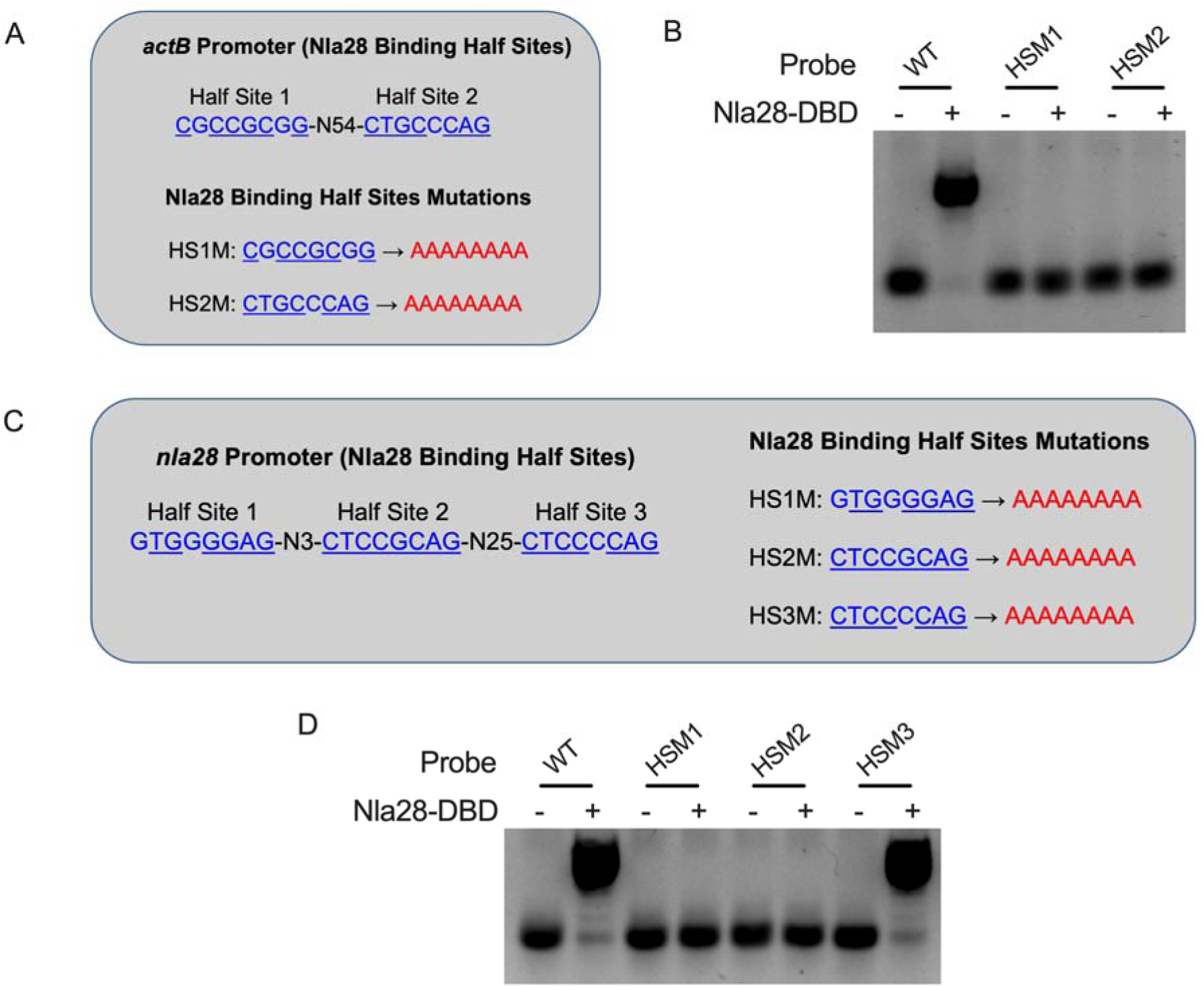
Two 8-bp sequences in the *actB* promoter and two in the *nla28* promoter are crucial for in vitro Nla28-DBD binding. Two 8-bp sequences (Half Site 1 and 2, shown in blue) that closely match the consensus Nla28 half binding were identified in the wild-type *actB* promoter fragment. Underlined sequences represent nucleotides that match consensus Nla28 half binding site. Mutations (HS1M and HS2M, shown in red) in these putative Nla28 half binding sites were generated for in vitro Nla28-DBD binding analysis. (B) EMSAs were performed with (+) or without (−) 2 μM of purified Nla28-DBD and a Cy5 5’ end-labeled *actB* promoter fragment containing two wild-type (WT) Nla28 half binding sites, a mutated half site one (HS1M) or a mutated half site two (HS2M). (C) Three 8-bp sequences (Half Site 1, 2 and 3, shown in blue) that closely match the consensus Nla28 half binding were identified in the wild-type *nla28* promoter fragment. Underlined sequences represent nucleotides that match consensus Nla28 half binding site. Mutations (HS1M, HS2M and HS3M, shown in red) in these putative Nla28 half binding sites were generated for in vitro Nla28-DBD binding analysis. (D) EMSAs were performed with (+) or without (−) 2 μM of purified Nla28-DBD and a Cy5 5’ end-labeled *nla2B* promoter fragment containing three wild-type (WT) Nla28 half binding sites, a mutated half site one (HS1M), a mutated half site two (HS2M) or a mutated half site three (HS3M).

In the next experiment we examined Nla28-DBD binding to the *nla28* promoter fragment, which contains three 8-bp sequences or potential Nla28-DBD half binding sites. Four *nla28* promoter fragments were generated for the in vitro binding assays. One of the *nla28* promoter fragments contained wild-type half sites 1-3 and each of the remaining *nla28* promoter fragments contained two wild-type half sites and one half site converted to all A nucleotides (see Figure 2C). The results of EMSAs performed with 2 μM of Nla28-DBD and 5’ Cy5-labeled wild-type and mutant *nla28* promoter fragments are shown in Figure 2D. Nla28-DBD binding was detected when the *nla28* promoter fragment contained all wild-type half sites, as previously noted. Nla28-DBD binding was also detected when the *nla28* promoter fragment contained wild-type half sites 1 and 2, and a half site 3 that was converted to all A nucleotides. However, no Nla28-DBD binding was detected when half site 1 or a half site 2 was converted to all A nucleotides, even though the remaining two half sites were wild type. Thus, it seems that half sites 1 and 2, but not half site 3 in the *nla28* promoter are crucial for in vitro Nla28-DBD binding. This is not the predicted result based solely on a putative half site’s similarity to the consensus, as half sites 1, 2 and 3 have 2, 0 and 1 mismatch(es), respectively, relative to the consensus Nla28 half binding site (Figure 2D and Table 1). Perhaps some nucleotide positions in a half site are more important for Nla28-DBD binding than others and/or the spacing between one half site and its neighboring half site is important for the efficient binding of a Nla28-DBD dimer.

In an additional experiment, the region of the *nla28* promoter containing half sites 1-3 was inserted into the *dev* promoter fragment, which is a fragment of a non-σ^54^ promoter and the negative control in previous EMSAs. As shown in Figure S2, a slower migrating and presumably Nla28-DBD-bound DNA complex was detected when 1.0 μM of Nla28-DBD was incubated with the ^32^P-labeld *dev* promoter fragment containing the inserted half sites, but not when 1.0 μM of Nla28-DBD was incubated with the ^32^P-labeld wild-type *dev* promoter fragment. These findings are consistent with the idea that the half sites we identified in the *nla28* promoter support in vitro Nla28-DBD binding.

### Nla28-DBD has relatively high affinity for the *nla28* promoter

The binding data presented here, in addition to previous binding and expression analyses (5, 21), suggest that Nla28 directly regulates the *nla28* operon in vivo. We speculated that Nla28 might have a relatively high affinity for its own promoter to produce a burst of *nla28* mRNA and presumably Nla28 protein early in development, which would in turn allow Nla28 to active transcription at downstream developmental promoters with low affinity binding sites. To examine this idea, we used various concentrations of purified Nla28-DBD in binding reactions with 5’ Cy5-labeled fragments of five putative target promoters, including the *nla28* promoter itself. As shown in Figure 3A-E, at a concentration of 0.5 μM we only detected Nla28-DBD binding to the *nla28* promoter, although we should note that binding to the MXAN2511 promoter was detected when Nla28-DBD was at the next lowest concentration of 0.75 μM. As for the remaining promoter fragments, we detected binding when Nla28-DBD was at a concentration of 1.0 μM (*actB*) or 1.5 μM (*pilA* and MXAN5040).

**Fig 3.**
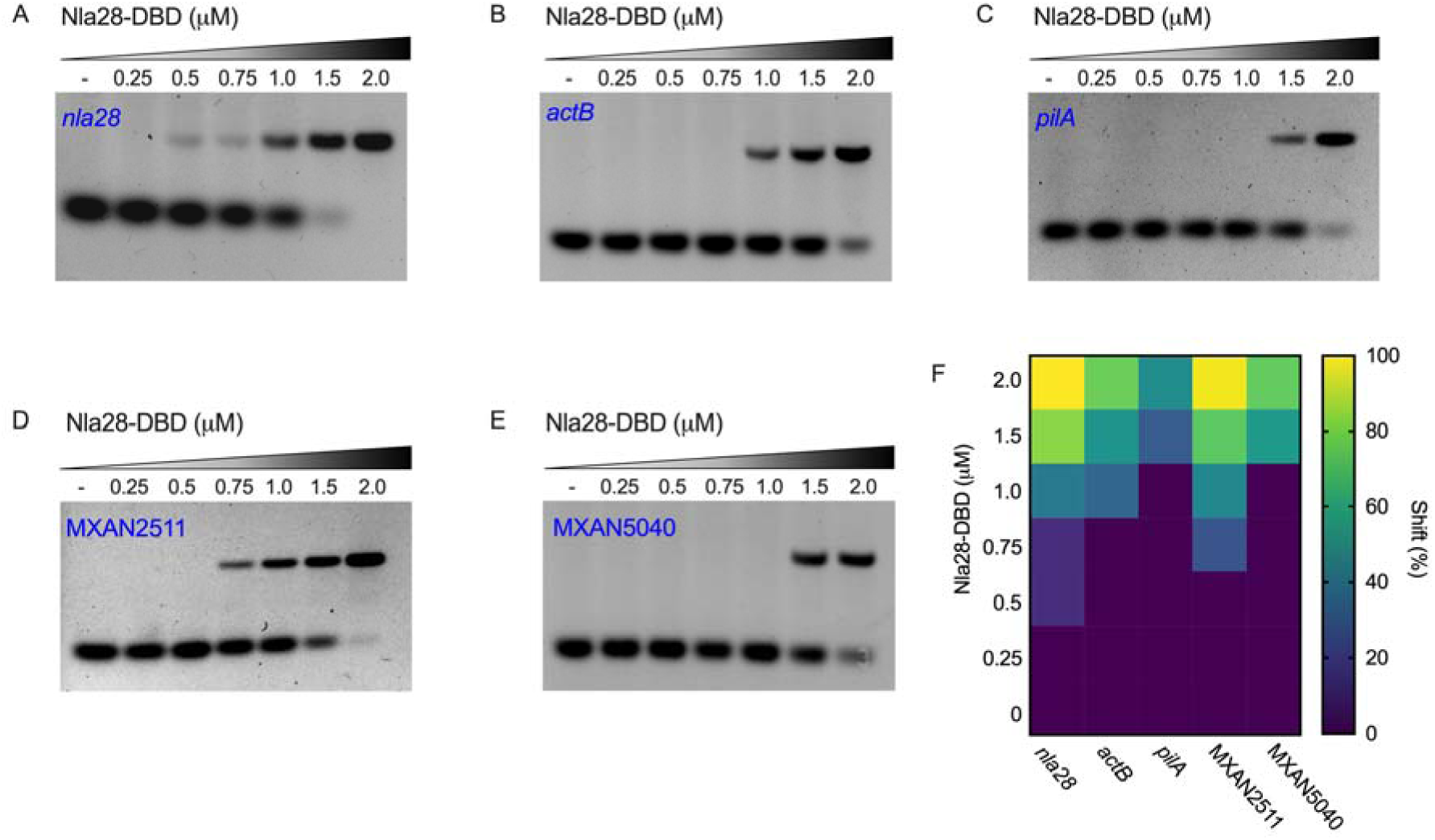
Nla28-DBD has relatively high affinity for the *nla28* promoter. The affinity of various concentrations of Nla28-DBD binding to targeted promoter DNA fragments were assessed by EMSAs. (A to E) Binding assays were performed with no purified Nla28-DBD added (-) or with Nla28-DBD added at the indicated concentration (0.25, 0.5, 0.75, 1.0, 1.5 or 2.0 μM) and a Cy5 5’ end-labeled fragment of the *actB* (A), *nla28* (B), *pilA* (C), MXAN2511 (D) or MXAN5040 (E) promoter in native PAGE. (F) Heat map representing color-coded quantitative evaluation on the affinity of various concentrations of Nla28-DBD binding to targeted promoter. Signal intensity in native PAGE was analyzed by Fiji ImageJ. Shift percentage (%) was calculated as follows: signal intensity shifted Nla28-DBD-bound DNA complex / (shifted Nla28-DBD-bound DNA complex + unshifted free DNA). Data are expressed as mean of triplicates from three independent experiments.

To confirm these findings, the above experiments were repeated two times and the signal intensities of shifted and unshifted complexes were determined (see *Materials and Methods*). Using these signal intensities, we then determined the average percentage of the total DNA that is in a shifted complex (ie., bound by Nla28-DBD) when each concentration of Nla28-DBD is mixed with each promoter fragment. The results, which are shown in Figure 3F, support the findings of the original titration series and that Nla28-DBD has a relatively high affinity for the *nla28* promoter, which likely facilitates autoregulation at low cellular concentrations. These results also support the proposed placement of the *nla28* promoter at or near the top of the hierarchy of Nla28-mediated transcription in developing cells.

### The developmental expression levels and patterns of genes located downstream of the promoters is altered in a *nla28* mutant

The results of the Nla28-DBD binding assays suggested that the 12 promoters listed in Table 1 might be in vivo targets of Nla28. Previous work indicated that peak developmental expression of *actB*, *nla6* and *nla28* is substantially reduced in a *nla28* mutant (5), which is consistent with this idea. To further test this idea, we examined whether a *nla28* mutation reduces developmental expression of genes located downstream of the newly identified promoter targets of Nla28 (see Table 1 and Figure S1). In particular, developmental expression of the single gene or one gene in the operon located downstream of the promoters were analyzed via quantitative PCR (qPCR); we examined the developmental mRNA levels of the genes in the *nla28* mutant and the corresponding wild-type strain. We should note that we selected the MXAN989 gene for qPCR analysis, even though the putative Nla28 target promoter is located within the MXAN989 gene. Our rationale in this case was that the predicted transcript from this promoter, which is located in the 5’ end of the MXAN989 gene, would contain the vast majority of the MXAN989 coding sequence and hence might yield a functional MXAN989 protein (Figure S1).

The relative mRNA levels in wild-type cells and *nla28* mutant cells at various developmental time points are shown in Figure 4. In wild-type cells, the mRNAs of *pilA* and MXAN5040 did not significantly increase during the developmental time course, a finding that contrasts with the results of a previous microarray analysis (5) and might be due to differences in the experimental methods and/or the developmental time points used in the studies. As for the remaining mRNAs analyzed via qPCR, significant increases were observed during the development of wild-type cells; peak developmental levels ranged from about 2.6 to 20.5-fold higher than the baseline vegetative (0 hours) levels. Furthermore, many of the mRNAs in the wild-type time course displayed an early (1 hour), pre-aggregation stage peak that was significantly above the vegetative baseline, and most of the mRNAs displayed a second, aggregation stage developmental peak (8-12 hours) that was significantly above the vegetative baseline. In the *nla28* mutant, the vegetive baseline levels of most mRNAs were reduced, suggesting that Nla28 modulates expression of these mRNAs during vegetative growth and supporting the previous assertion that Nla28 is not a fruiting body development-specific transcriptional activator (21). As for mRNA levels in developing *nla28* mutant cells, the peaks were substantially lower than in wild-type cells or the peaks observed in wild-type cells were absent. These findings indicate that Nla28 also modulates expression of these mRNAs during development.

**Fig 4.**
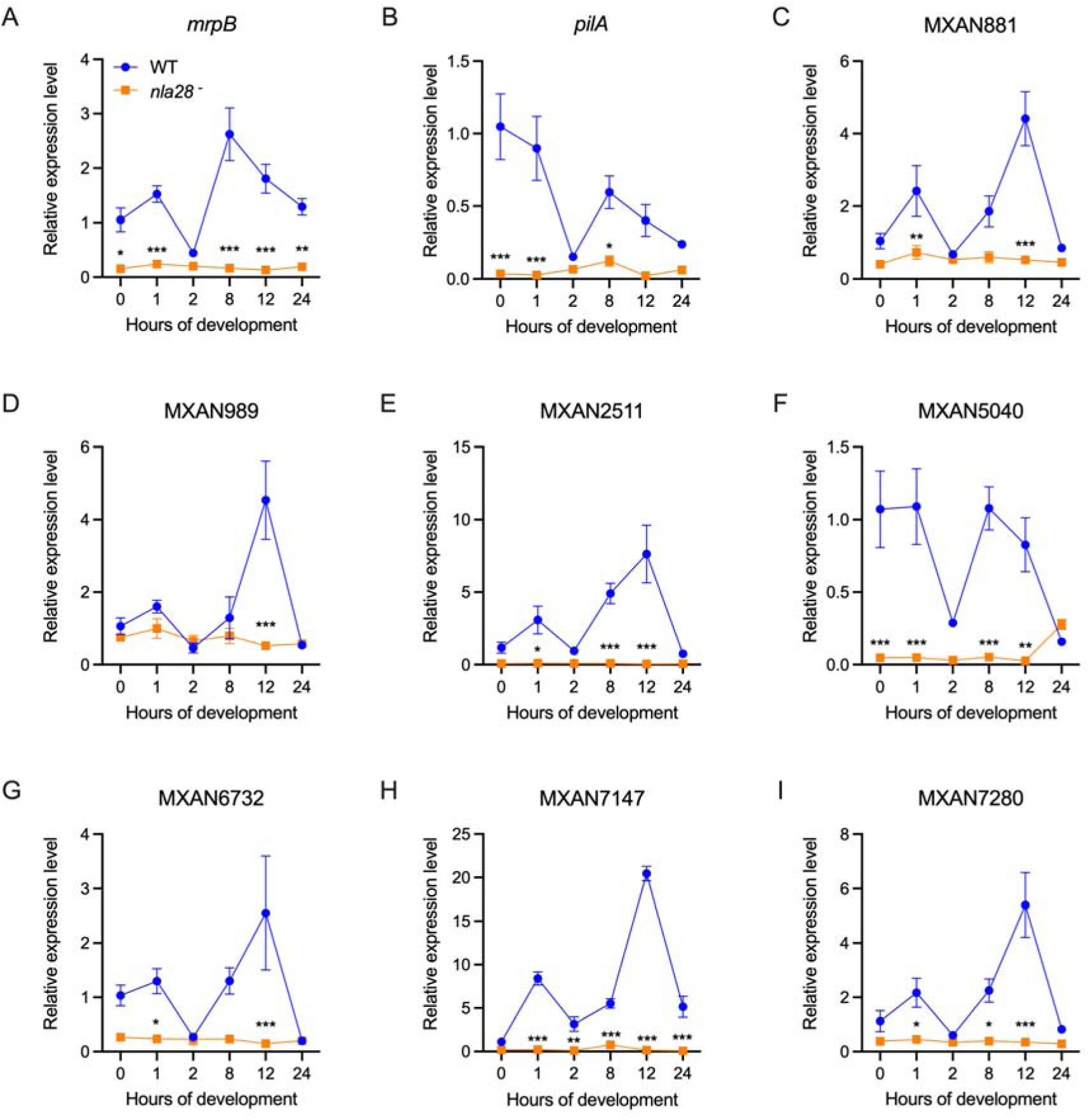
The developmental expression patterns of Nla28 target genes are altered in a *nla28* mutant. The developmental mRNA levels of the *mrpB* (A), p*ilA* (B), MXAN881 (C), MXAN989 (D), MXAN2511 (E), MXAN5040 (F), MXAN6732 (G), MXAN7147 (H) and MXAN7280 (I) genes in the *nla28* mutant (*nla28*^-^) and the corresponding wild-type(WT) strain were determined using qPCR. Wild-type and *nla28* mutant cells were harvested at 0, 1, 2, 8, 12 and 24 hours of development for RNA isolation and qPCR analysis. N = 3 technical replicates of pooled RNA samples at each time point. Error bars are standard deviations of the means. The data was analyzed using two-way analysis of variance (ANOVA) and Tukey’s multiple comparisons *post hoc* tests; ***p < 0.001; **p < 0.01; *p < 0.05 for mRNA levels in *nla28* mutant versus wild-type cells.

### Mutations in the putative Nla28 binding site substantially reduce *nla28* promoter activity in developing cells

The in vitro binding and in vivo expression data presented here and in previous work (5) suggested that Nla28 might directly regulate the 12 target promoters we identified. To further examine this idea, we selected the *nla28* promoter for an in vivo mutational analysis. We started with a fragment of the *nla28* promoter region that contains the putative σ^54^-RNA polymerase and Nla28 binding sites and generated fragments with a 2-bp substitution, 4-bp substitution or 6-bp substitution in Nla28 half site 2. Wild-type and mutant *nla28* promoter fragments were cloned into pTL1, a plasmid that creates transcriptional fusions between cloned promoters and the *lacZ* gene. The *lacZ* fusion plasmids were introduced into wild-type strain DK1622, and Kan^r^ isolates carrying a plasmid integrated (via a single homologous recombination event) into the chromosomal *nla28* locus were identified via PCR and analyzed via DNA sequencing. All isolates contained an upstream copy of the *nla28* promoter fused to *lacZ*, followed by plasmid DNA, and then a second copy of the *nla28* promoter followed by an intact *nla28* operon. Isolates with two wild-type copies of the *nla28* promoter (i.e., the wild-type strains for these experiments), or an upstream mutant copy and a wild-type downstream copy of the *nla28* promoter (i.e., Nla28 half site mutants) were allowed to develop for various amounts of time and β-Galactosidase assays were used to infer developmental promoter activities.

As shown in Figure 5, all the half site 2 mutations reduced the peak in vivo activity of the *nla28* promoter during development; the peak activities of the mutant *nla28* promoters ranged from about 1.5- to 2.4-fold less than wild-type peak activity. An additional experiment was performed with two derivatives of plasmid pREG1727; pREG1727 confers kanamycin resistance (Kan^r^) and carries the Mx*8 attP* site, which allows site-specific integration at the Mx*8* phage attachment site in the *M*. *xanthus* chromosome (26). In particular, the wild-type *nla28* promoter fragment and the *nla28* promoter fragment carrying the 2-bp substitution in Nla28 half site 2 were cloned into pREG1727, the plasmids were introduced into wild-type strain DK1622, Kan^r^ isolates carrying a plasmid integrated at the ectopic Mx*8* phage attachment site were identified via PCR and DNA sequencing, in vivo developmental promoter activities were monitored as noted above, and the activities were similar to their counterparts integrated at the native *nla28* locus (data not shown).

**Fig 5.**
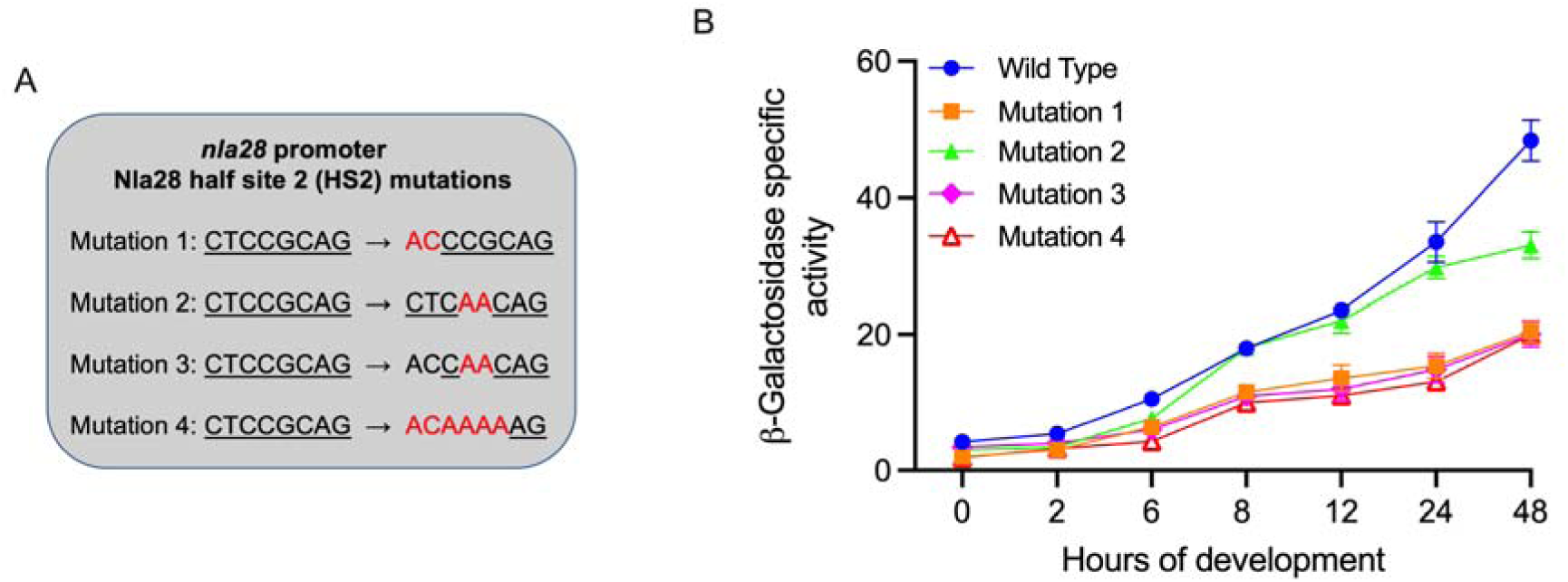
Mutations in the putative Nla28 binding site substantially reduce *nla28* promoter activity in developing cells. (A) *nla28* promoter fragments with a 2-bp substitution (Mutation 1 and 2), 4-bp substitution (Mutation 3) or 6-bp substitution (Mutation 4) in Nla28 half site 2 were generated by site-directed mutagenesis. Substituted nucleotides are shown in red. (B) Wild-type *nla28* promoter and derivatives of the *nla28* promoter carrying Mutation 1, Mutation 2, Mutation 3 or Mutation 4 were cloned into a *lacZ* expression vector and transferred to the wild-type *M. xanthus* strain DK1622. In vivo activities of wild-type and mutant *nla28* promoters were inferred from β galactosidase-specific activities (defined as nanomoles of ONP produced per minute per milligram of protein) at various time points (0, 2, 6, 8, 12, 24 and 48 h) during development. N = 3 biological replicates at each of the indicated time points. Error bars are standard deviations of the means.

We should note that the developmental activity of the *nla28* promoter is modulated by the Nla28 and Nla6 EBPs (5, 18) and we were not expecting *nla28* promoter activity to be abolished when we mutated half site 2. Indeed, the developmental activity of the *nla28* promoter is reduced about 2.6-fold when the *nla28* gene is inactivated, but developmental activity is not abolished (5). Thus, the Nla28 half binding site that we identified using in vitro assays is important for in the vivo developmental activity of the *nla28* promoter and it seems likely that the Nla28 directly regulates the *nla28* promoter in vivo.

### Nla28 target genes are important for development

The results of our in vitro and in vivo expression studies indicate that the loci shown in Table 1 are likely to be in vivo targets of Nla28 during development. Four of the loci (*actB*, *nla6*, *nla28* and *mrpB*) contain characterized EBP genes and the corresponding EBPs are known to be important for *M*. *xanthus* development (14, 18, 33–35). Another locus that has been characterized is *pilA*; this locus is known to be important for production of type IV pili, social motility and normal development (36–38).

To examine whether the remaining Nla28 targets listed in Table 1 are important for development, we generated an insertion in one gene in each locus. Namely, we generated an insertion in the single gene or in one gene in the operon located downstream of the putative core σ^54^ promoter element. Subsequently, we determined whether the insertions affect the formation of aggregates of cells and/or sporulation. The results of these phenotypic studies are summarized in Table 2. Briefly, all the insertions affected the timing of aggregate formation and some also affected the final shape of aggregates; however, none of the insertions completely blocked the formation of aggregates. In addition, the insertions substantially reduced the number of sonication- and heat-resistant spores that were able to germinate into colonies, as indicated by the viable spore numbers shown in Table 2. Interestingly, the insertions had relatively minor impacts on spore numbers; spore number refers to the number of developing cells that were able make the shape change (rod to spherical shape) associated with the early differentiation stage of sporulation. Thus, it seems that the insertions had a stronger impact on the maturation of spores into stress resistance cells and/or on spore germination than on spore differentiation.

**TABLE 2.** Developmental phenotypes of wild-type and mutant cells

| Strain | Aggregation<sup>a</sup> | Fruiting body sporulation<sup>b</sup> |  |
| --- | --- | --- | --- |
|  |  | Spore number (%) | Viable spore number (%) |
| DK1622 | + | 100±12.6 | 100±8.5 |
| *nla28* | +/- | 108.1±7.6 | 2.8±1.2 |
| MXAN0881 | +/- | 72.2±0.7 | 0.5±0.5 |
| MXAN0989 | +/- | 91.1±5.0 | <0.1 |
| MXAN5040 | +/- | 55.5±7.3 | 1.2±0.8 |
| MXAN7147 | +/- | 78.5±8.1 | 7.9±2.5 |
| MXAN7279 | +/-, - | 44.0±7.8 | 5.4±2.1 |
| MXAN2510 | +/-, - | 35.0±8.6 | <0.1 |
| MXAN6732 | - | 47.4±3.7 | 6.3±3.3 |

<sup>a</sup> +, produced normal-looking aggregates; +/-, aggregated failed to compact or darken as quickly as wild-type aggregates; -, aggregates (after 5 days of development) had abnormal shapes.
<sup>b</sup>The mean values (± standard deviations) for the spore assays are shown as percentages of *M. xanthus* DK1622 (wild type). The number of fruiting body spores produced by wild-type cells ranged from $6 \times 10^6$ to $4 \times 10^7$ . Means (± standard deviations) derived from three independent experiments are shown.

### Insertions in some Nla28 target genes affect swarm expansion

As noted above, one of the characterized targets of Nla28 is *pilA*, a gene that is important for type IV pili-based social motility, surface spreading and normal development (36–38). Since many of the remaining Nla28 targets have yet to be tested for roles in *M*. *xanthus* motility, we examined whether mutations in Nla28 target genes affect colony spreading on agar surfaces. Specifically, cells carrying mutations in Nla28 target genes were placed on the surface of 1.5% and 0.4% agar plates, the plates were incubated at 32°C for 3 days, and the colony diameters were compared to those produced by wild-type cells and the negative control cells, which are unable to actively spread on agar surfaces due to mutations that inhibit social motility and adventurous motility (*i.e*., the two motility systems that *M*. *xanthus* uses for surface spreading) (39, 40).

The results shown in Table 3 indicate that most of the mutations in Nla28 target genes have little or no impact on surface spreading. Insertions in MXAN2510 and MXAN7279 are notable exceptions, as strains carrying an insertion in either gene show substantial reductions in surface spreading compared to the wild-type strain. In the case of the MXAN7279 insertion mutant, a reduction in surface spreading was observed on 0.4% agar plates, which provide a soft and wet surface that favors social motility, but not on 1.5% agar plates, which provide a relatively firm and dry surface that favors A-motility (41). As for the MXAN2510 insertion mutant, surface spreading on 1.5% agar plates was slightly reduced and surface spreading on 0.4% agar plates was substantially reduced. This phenotype is reminiscent of the *pilA* (social motility) mutant, which displays reduced surface spreading on both agar surfaces, but the reduction is most dramatic on 0.4% agar.

**TABLE 3.** Swarm diameters of wild-type and mutant cells on 0.4% and 1.5% agar^a^

| Strain | Mean swarm diameter (percentage of wild type) |  |
| --- | --- | --- |
|  | Soft agar (0.4%) | Hard agar (1.5%) |
| DK1622 (wild type) | 100 ± 10 | 100 ± 5 |
| DK 2161 (A<sup>-</sup> S<sup>-</sup>)<sup>b</sup> | 38 ± 3 | 37 ± 3 |
| *mrpB* | 94 ± 8 | 100 ± 5 |
| *nla28* | 96 ± 5 | 93 ± 6 |
| *pilA* | 27 ± 10 | 72 ± 6 |
| MXAN0881 | 101 ± 10 | 96 ± 11 |
| MXAN0989 | 88 ± 11 | 81 ± 10 |
| MXAN2510 | 48 ± 3 | 82 ± 7 |
| MXAN5040 | 89 ± 5 | 82 ± 4 |
| MXAN6732 | 108 ± 12 | 108 ± 4 |
| MXAN7147 | 96 ± 7 | 94 ± 4 |
| MXAN7279 | 41 ± 7 | 110 ± 3 |

<sup>a</sup>The mean diameter (± standard deviation) of five swarms produced by each mutant strain was determined and normalized to the mean diameter of five swarms produced by wild-type strain DK1622.
<sup>b</sup>A<sup>-</sup> represents defect in adventurous motility (A-motility); S<sup>-</sup> represents defect in social motility (S-motility).

## DISCUSSION

### The likely binding site of Nla28 is an 8-bp direct repeat sequence

One goal of this study was to better understand how Nla28 identifies its target developmental promoters. Using a collection of developmental promoter targets (Table 1), in vitro promoter binding assays (Figure 1) and in vitro mutational analyses (Figures 3 and 4), we were able to identify tandem (imperfect) repeats of an 8-bp sequence [consensus CT(C/G)CG(C/G)AG] as the probable binding site of Nla28 dimers. Our in vivo studies further suggest that the tandem repeat sequences, as well as the Nla28 protein, are important for function of the target promoters in developing *M*. *xanthus* cells. In many of these promoters, the repeats or Nla28 half binding sites are found in clusters (Figure S1). In fact, seven promoters have multiple tandem repeats that closely match the consensus; we scanned the 300 bp of DNA upstream of the -12- and -24-bp regions for such repeats. Thus, most of the promoters have multiple sites for potential binding of Nla28 dimers. One possible explanation for this arrangement is that the clustering of binding sites helps sequester Nla28 dimers at the promoters, increasing the local concentrations. A relatively high local concentration of Nla28 would in turn facilitate transcription, even when the individual binding sites in the promoter are low affinity sites (42–44). Indeed, this might be the situation in the *pilA* and MXAN5040 promoters, as these promoters have multiple sites for the binding of Nla28 dimers and the sites that have been tested for Nla28-DBD binding appear to be relatively low affinity sites (Figure 3).

The *nla28* promoter seems to be on the opposite end of the spectrum; this promoter contains three potential half sites (Table 1), only two of the half sites are important for Nla28-DBD binding (Figure 2D) and a Nla28 dimer is likely to have a high affinity for these two half sites (Figure 3). As previously suggested, Nla28 might use a relatively high affinity binding site in its own promoter to produce a development-associated burst of *nla28* mRNA and Nla28 protein. The higher levels of Nla28 would in turn facilitate transcription at developmental promoters containing low affinity binding sites. We should note that the increase in *nla28* expression in developing cells might be connected to the presence of A-signal, which is a starvation-induced cell density signal (11–13), via the Nla28S/Nla28 signal transduction system, as previously proposed (19, 20).

### Potential Nla28 binding sites are found upstream of many developmental genes

Since we were interested in getting a more global view of the numbers and types of developmental genes that Nla28 directly regulates, we used the *M*. *xanthus* genome sequence, the consensus Nla28 half binding site, bioinformatics and expression data to identify additional promoter/gene targets of Nla28 (see *Materials and Methods*). Briefly, we looked for tandem 8-bp sequences with no more than two mismatches relative to the consensus Nla28 half binding site and putative σ^54^-RNA polymerase binding sites located downstream of these tandem sequences (Table S3). We also identified single genes and operons located downstream of the putative σ^54^-RNA polymerase binding sites, and used DNA microarrays to determine whether the downstream gene or genes are developmentally regulated and whether normal expression in developing cells is dependent on Nla28 (5). As shown in Table S4, an additional 58 putative σ promoters and 102 genes were tagged as potential targets of Nla28 using the consensus Nla28 binding sequence, bringing the total number of candidates for direct Nla28 regulation to 70 promoters and 140 genes. Interestingly, many of the putative promoters that we identified have multiple sites for Nla28 dimer binding, which is consistent with the initial 12 promoters that we examined. The types of genes that were discovered in this analysis are discussed below.

### The vast majority of Nla28 target promoters are intragenic

In a recent study of natural product gene regulation in *M*. *xanthus* (21), the majority of experimentally confirmed and putative σ^54^-RNA polymerase and Nla28 binding sites were localized to natural product genes and not to intergenic sequences. In the analysis of developmental promoter targets of Nla28 described here, we obtained similar results (summarized in Figure S1). With these findings and previous data linking the majority of the σ^54^ promoter targets of Nla6 to intragenic regions (18), a pattern of intragenic σ^54^ promoter usage in *M*. *xanthus* is starting to emerge. However, we are still unable to make sweeping conclusions regarding the genomic locations of *M*. *xanthus* promoters, as large-scale promoter analyses are lacking. This is especially true for σ^70^-type promoters, which are more common than σ^54^ promoters (45), but a notable study of the binding sites of the σ^70^-type transcription factor MrpC was published a few years ago. Specifically, a ChIP-seq analysis of the genomic binding sites of the MrpC protein yielded many potential intragenic locations; 61% of the ChIP-seq peaks were in coding regions (46).

Although genome-wide promoter analyses are generally lacking in other bacterial species, potential genomic binding sites for σ^54^ have been identified in some well-studies systems. For example, ChIP-chip analyses in *Salmonella enterica* Serovar Typhimurium linked the majority of potential σ^54^ binding sites to intragenic regions (47, 48). In *Escherichia coli*, one ChIP-seq analysis placed 2/3 of putative σ^54^ binding sites in intragenic regions (49). While some of the intragenic σ^54^ binding sites such as those identified in *Salmonella* have been confirmed using other methods (ie., EMSAs) (48), providing more confidence in the results, the bulk of the intragenic sites await such conformation. Hence, the results of the above studies are intriguing, but the question of how commonly σ^54^-RNA polymerase uses intragenic sites in *Salmonella* and *E*. *coli*, not to mention other bacterial species, remains open.

The question of how often bacteria use intragenic σ^70^-type promoters also remains open. A notable study attempted to address this question using ChIP-seq and RNA-seq (50). This study, which focused on the σ^70^-type sigma factor FliA from *E*. *coli*, linked many of the genomic FliA binding sites to intragenic regions. However, further analyses revealed that the vast majority of the intragenic promoters yielded unstable RNAs or were inactive under the selected experimental conditions, leading the authors to conclude that most of the intragenic promoters are not important *cis*-regulatory elements (51).

Finally, it is worth reminding the reader that in vivo mutational analyses have been performed on several intragenic promoter targets of Nla6 and Nla28 (5, 21, 29) and the results indicate that the promoters have the signature properties of σ^54^ promoter elements and are crucial for developmental and/or growth-phase related gene expression. Thus, it seems these intragenic targets of Nla6 and Nla28 are bona fide σ^54^ promoters, as predicted. Given the experimental confirmation of these intragenic promoters, we suggest that having a promoter in the coding sequences of genes might have advantages in some cases. For example, some of these putative σ^54^ promoters are in the coding sequence of a gene that is far upstream of the single gene or operon they are predicted to regulate. This distant location would produce a relatively long 5’ untranslated region (UTR) in the mRNA and perhaps the 5’ UTR provides an additional layer of regulation for natural product or developmental gene expression. Other intragenic σ^54^ promoters appear to be internal promoters; promoters located within a gene in an operon instead of upstream of the first gene in the operon. In such cases, the promoter is predicted to yield an mRNA corresponding to a subset of an operon’s genes, providing flexibility if only some of the genes are needed in a particular environment (52).

### Most of the confirmed Nla28 targets have regulatory functions or are predicted to have defense functions

As noted above, the molecular/cellular and developmental functions of many of the confirmed Nla28 targets were identified prior to this work. Regulatory or signal transduction functions dominate this group of Nla28 targets, as all but one of these loci fall into this category. This includes the *actB*, *nla6*, *nla28* and *mrpB* loci, which contain genes that code for EBPs; these EBPs modulate transcription of different types of developmental genes (14, 15, 33–35, 53–55); also, see Tables 1 and S4. The EBP loci also contain genes for signal transduction proteins such as the histidine protein kinase partners of Nla6 and Nla28, which are called Nla6S and Nla28S, respectively (15, 18–20, 32). The one confirmed and previously characterized Nla28 target that does not have a regulatory function is *pilA*; the *pilA* locus contains genes that are important for type IV pili-based social motility, surface spreading and normal development (14, 15, 36–38, 56, 57).

All confirmed Nla28 targets characterized in this study are important for development and two are also important for surface spreading and presumably motility (Tables 2 and 3). The most common molecular/cellular functional category assigned to these loci, based on the annotated genome sequence of wild-type strain DK1622 (15), is defense mechanisms (Table S5 and Figure 6A). This includes genes that are predicted to encode heavy metal efflux transporter components. It also includes CRISPR-associated proteins, which have been characterized in a variety of bacterial species and are known to be involved in adaptive immunity against foreign genetic elements (58–62). As for the remaining Nla28 target loci characterized here, two have been linked to potential metabolic functions. Namely, one locus is predicted to be involved in lipid metabolism and the second in carbohydrate metabolism. The other two loci characterized here have not been assigned a particular function. However, the three Nla28 targets characterized in a recent study (5, 20, 21) are worth mentioning here, since they are likely involved in production of secondary metabolites and hence could have a defense function or could be involved in signaling.

**Fig 6.**
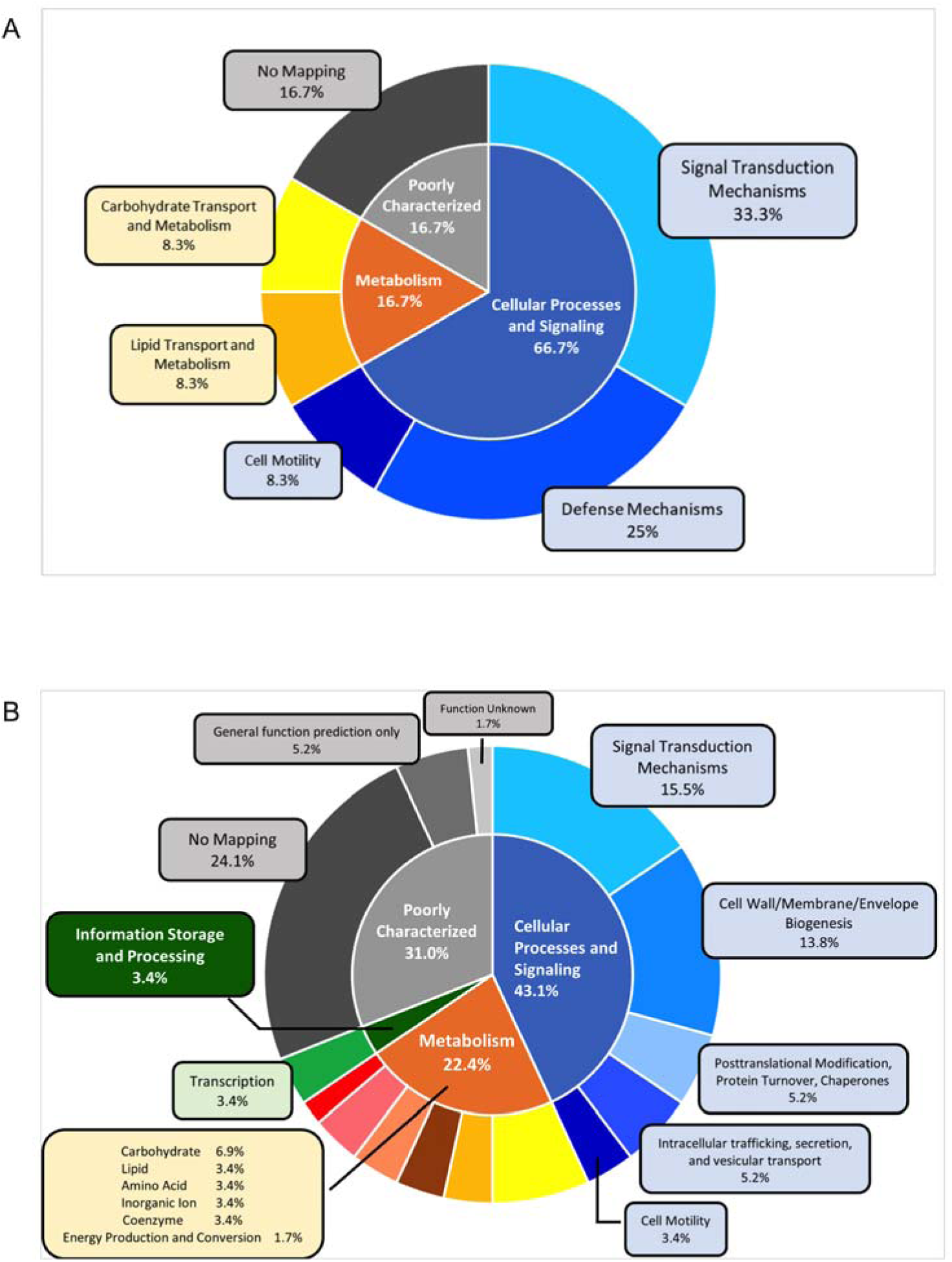
Functional category and subcategory distributions of Nla28 targets based on the Clusters of Orthologous Groups of proteins (COGs) database. The pie chart shows the proportion of confirmed Nla28 targets with known/unknown molecular/cellular functions: 66.7% are associated with cellular processes and signaling (33.3% signal transduction mechanisms; 25% defense mechanisms; 8.3% cell motility); 16.7% are associated with metabolism (8.3% carbohydrate transport and metabolism; 8.3% lipid transport and metabolism); 16.7% are poorly characterized (16.7% no mapping). (B) The pie chart shows the proportion of putative Nla28 targets with known/unknown molecular/cellular functions: 43.1% are associated with cellular processes and signaling; 22.4% are associated with metabolism; 3.4% are associated with information storage and processing; 31% are poorly characterized. Various subcategories are also shown.

It has been suggested that Nla28 is part of a general stress response induced by nutrient depletion, since Nla28 modulates transcription of many early, starvation-responsive developmental genes and genes that are highly expressed during the transition into stationary phase (5, 20, 21). That some confirmed targets have defense functions makes sense if Nla28 is viewed as a regulator of starvation-induced stress. The metabolic functions of other Nla28 targets also make sense in this context, as one would expect nutrient depletion to be accompanied by changes in cellular metabolism. Additional information that supports the view of Nla28 as a regulator of *M*. *xanthus*’ stress response comes from physiological studies in other Gram-negative bacteria. In particular, orthologues of MXAN881, MXAN989, MXAN5040, MXAN7147 and MXAN7280 have all been linked to stress responses in other Gram-negative bacteria (63–67).

### Regulatory, metabolic and cell envelope biogenesis are common functions among putative Nla28 targets

As noted above, 58 putative σ^54^ promoters and 102 genes were tagged as potential targets of Nla28 using the consensus Nla28 half binding site, bringing the total number of candidates for direct Nla28 regulation to 70 promoters and 140 genes. Based on the annotated genome sequence of wild-type strain DK1622 (15), we were able to place the putative Nla28 targets into functional molecular/cellular functional categories (Table S6).

Figure 6B shows the percentages of putative Nla28 targets that were placed into each category. The three most highly represented categories are metabolic functions, regulatory/signal transduction and cell envelope/cell wall biogenesis. Given that developing cells are experiencing starvation, changes in cellular metabolism might be expected. Similarly, developing cells might be expected to express genes involved in cell wall or cell membrane structural changes, as developing cells are experiencing starvation-induced stress. Other notable categories of putative Nla28 targets include motility genes, post-translational modifications/protein turnover and protein secretion. Finally, many of the putative Nla28 targets have predicted orthologues linked to stress responses in other Gram-negative bacteria. For example, potential orthologues of MXAN0162, MXAN0934 MXAN2907 have been linked to envelope stress in Gram – species (68–71).

### Conclusions

Nla28 is an EBP that begins modulating gene expression soon after *M*. *xanthus* cells encounter nutrient-poor conditions. In this study, we identified the direct targets of Nla28 to better understand Nla28’s function and to help define the early gene regulatory pathways involved in *M*. *xanthus*’ starvation response. We found that Nla28 dimers recognize their target promoters using tandem, imperfect repeats of an 8-bp sequence and that most of the Nla28 target promoters are intragenic. Seventy promoters and 140 genes were classified as potential targets of Nla28, suggesting that the Nla28 regulon might be relatively large. Common functions assigned to Nla28 target genes include regulatory, metabolic and defense-related functions, and cell envelope biogenesis. Many of these functions make sense in the context of Nla28’s role as a general regulator of stress-associated genes and, based on the work presented here, we now know that several these genes are important for production of stress-resistant spores following starvation.

## Abbreviations

EBP: enhancer binding protein
DBD: DNA binding domain

## REFERENCES

1. Rajagopalan R, Kroos L. 2014. Nutrient-regulated proteolysis of MrpC halts expression of genes important for commitment to sporulation during Myxococcus xanthus development. Journal of bacteriology 196:2736–2747.

2. Popham DL, Szeto D, Keener J, Kustu S. 1989. Function of a bacterial activator protein that binds to transcriptional enhancers. Science 243:629–635.

3. Sasse-Dwight S, Gralia JD. 1990. Role of eukaryotic-type functional domains found in the prokaryotic enhancer receptor factor σ54. Cell 62:945–954.

4. Wedel A, Kustu S. 1995. The bacterial enhancer-binding protein NTRC is a molecular machine: ATP hydrolysis is coupled to transcriptional activation. Genes & development 9:2042–2052.

5. Giglio KM, Caberoy N, Suen G, Kaiser D, Garza AG. 2011. A cascade of coregulating enhancer binding proteins initiates and propagates a multicellular developmental program. Proceedings of the National Academy of Sciences 108:E431–E439.

6. O’Connor KA, Zusman DR. 1991. Development in Myxococcus xanthus involves differentiation into two cell types, peripheral rods and spores. Journal of Bacteriology 173:3318–3333.

7. O’Connor KA, Zusman DR. 1991. Behavior of peripheral rods and their role in the life cycle of Myxococcus xanthus. Journal of bacteriology 173:3342–3355.

8. O’Connor KA, Zusman DR. 1989. Patterns of cellular interactions during fruiting-body formation in Myxococcus xanthus. Journal of bacteriology 171:6013–6024.

9. Singer M, Kaiser D. 1995. Ectopic production of guanosine penta-and tetraphosphate can initiate early developmental gene expression in Myxococcus xanthus. Genes & development 9:1633–1644.

10. Harris BZ, Kaiser D, Singer M. 1998. The guanosine nucleotide (p) ppGpp initiates development and A-factor production inMyxococcus xanthus. Genes & development 12:1022–1035.

11. Kuspa A, Plamann L, Kaiser D. 1992. A-signalling and the cell density requirement for Myxococcus xanthus development. Journal of Bacteriology 174:7360–7369.

12. Kuspa A, Plamann L, Kaiser D. 1992. Identification of heat-stable A-factor from Myxococcus xanthus. Journal of Bacteriology 174:3319–3326.

13. Plamann L, Kuspa A, Kaiser D. 1992. Proteins that rescue A-signal-defective mutants of Myxococcus xanthus. Journal of bacteriology 174:3311–3318.

14. Caberoy NB, Welch RD, Jakobsen JS, Slater SC, Garza AG. 2003. Global mutational analysis of NtrC-like activators in Myxococcus xanthus: identifying activator mutants defective for motility and fruiting body development. Journal of Bacteriology 185:6083–6094.

15. Goldman B, Nierman W, Kaiser D, Slater S, Durkin AS, Eisen JA, Ronning C, Barbazuk W, Blanchard M, Field C. 2006. Evolution of sensory complexity recorded in a myxobacterial genome. Proceedings of the National Academy of Sciences 103:15200–15205.

16. Diodati ME, Ossa F, Caberoy NB, Jose IR, Hiraiwa W, Igo MM, Singer M, Garza AG. 2006. Nla18, a key regulatory protein required for normal growth and development of Myxococcus xanthus. Journal of bacteriology 188:1733–1743.

17. Ossa F, Diodati ME, Caberoy NB, Giglio KM, Edmonds M, Singer M, Garza AG. 2007. The Myxococcus xanthus Nla4 protein is important for expression of stringent response-associated genes, ppGpp accumulation, and fruiting body development. Journal of bacteriology 189:8474–8483.

18. Giglio KM, Zhu C, Klunder C, Kummer S, Garza AG. 2015. The enhancer binding protein Nla6 regulates developmental genes that are important for Myxococcus xanthus sporulation. Journal of bacteriology 197:1276–1287.

19. Sarwar Z, Garza AG. 2012. The Nla28S/Nla28 two-component signal transduction system regulates sporulation in Myxococcus xanthus. Journal of bacteriology 194:4698–4708.

20. Sarwar Z, Garza AG. 2016. Two-component signal transduction systems that regulate the temporal and spatial expression of Myxococcus xanthus sporulation genes. Journal of bacteriology 198:377–385.

21. Ma M, Welch RD, Garza AG. 2021. The σ 54 system directly regulates bacterial natural product genes. Scientific reports 11:1–11.

22. Bush M, Dixon R. 2012. The role of bacterial enhancer binding proteins as specialized activators of σ54-dependent transcription. Microbiology and Molecular Biology Reviews 76:497–529.

23. Gao F, Danson AE, Ye F, Jovanovic M, Buck M, Zhang X. 2020. Bacterial Enhancer Binding Proteins—AAA+ Proteins in Transcription Activation. Biomolecules 10:351.

24. Giglio KM, Eisenstatt J, Garza AG. 2010. Identification of enhancer binding proteins important for Myxococcus xanthus development. Journal of bacteriology 192:360–364.

25. Plamann L, Davis JM, Cantwell B, Mayor J. 1994. Evidence that asgB encodes a DNA-binding protein essential for growth and development of Myxococcus xanthus. Journal of bacteriology 176:2013–2020.

26. Fisseha M, Gloudemans M, Gill RE, Kroos L. 1996. Characterization of the regulatory region of a cell interaction-dependent gene in Myxococcus xanthus. Journal of Bacteriology 178:2539–2550.

27. Kaplan H, Kuspa A, Kaiser D. 1991. Suppressors that permit A-signal-independent developmental gene expression in Myxococcus xanthus. Journal of bacteriology 173:1460–1470.

28. Wu SS, Kaiser D. 1997. Regulation of expression of the pilA gene in Myxococcus xanthus. Journal of bacteriology 179:7748–7758.

29. Gronewold TM, Kaiser D. 2007. Mutations of the act promoter in Myxococcus xanthus. Journal of bacteriology 189:1836–1844.

30. Studholme DJ, Buck M, Nixon T. 2000. Identification of potential σN-dependent promoters in bacterial genomes. Microbiology 146:3021–3023.

31. Schindelin J, Arganda-Carreras I, Frise E, Kaynig V, Longair M, Pietzsch T, Preibisch S, Rueden C, Saalfeld S, Schmid B. 2012. Fiji: an open-source platform for biological-image analysis. Nature methods 9:676–682.

32. Sarwar Z, Garza AG. 2012. The Nla6S protein of Myxococcus xanthus is the prototype for a new family of bacterial histidine kinases. FEMS microbiology letters 335:86–94.

33. Gorski L, Kaiser D. 1998. Targeted mutagenesis of ς54 activator proteins in Myxococcus xanthus. Journal of Bacteriology 180:5896–5905.

34. Gronewold TM, Kaiser D. 2001. The act operon controls the level and time of Cusignal production for Myxococcus xanthus development. Molecular microbiology 40:744–756.

35. Sun H, Shi W. 2001. Genetic studies of mrp, a locus essential for cellular aggregation and sporulation of Myxococcus xanthus. Journal of bacteriology 183:4786–4795.

36. Wu SS, Kaiser D. 1995. Genetic and functional evidence that type IV pili are required for social gliding motility in Myxococcus xanthus. Molecular microbiology 18:547–558.

37. Wu SS, Kaiser D. 1996. Markerless deletions of pil genes in Myxococcus xanthus generated by counterselection with the Bacillus subtilis sacB gene. Journal of Bacteriology 178:5817–5821.

38. Wu SS. 1998. The role of Type IV pili in social gliding motility of Myxococcus xanthus. Stanford University.

39. Hodgkin J, Kaiser D. 1979. Genetics of gliding motility in Myxococcus xanthus (Myxobacterales): two gene systems control movement. Molecular and General Genetics MGG 171:177–191.

40. Hodgkin J, Kaiser D. 1979. Genetics of gliding motility in Myxococcus xanthus (Myxobacterales): genes controlling movement of single cells. Molecular and General Genetics MGG 171:167–176.

41. Shi W, Zusman DR. 1993. The two motility systems of Myxococcus xanthus show different selective advantages on various surfaces. Proceedings of the National Academy of Sciences 90:3378–3382.

42. Sharon E, Kalma Y, Sharp A, Raveh-Sadka T, Levo M, Zeevi D, Keren L, Yakhini Z, Weinberger A, Segal E. 2012. Inferring gene regulatory logic from high-throughput measurements of thousands of systematically designed promoters. Nature biotechnology 30:521–530.

43. Ezer D, Zabet NR, Adryan B. 2014. Physical constraints determine the logic of bacterial promoter architectures. Nucleic acids research 42:4196–4207.

44. Ezer D, Zabet NR, Adryan B. 2014. Homotypic clusters of transcription factor binding sites: A model system for understanding the physical mechanics of gene expression. Computational and structural biotechnology journal 10:63–69.

45. Barrios H, Valderrama B, Morett E. 1999. Compilation and analysis of sigma(54)-dependent promoter sequences. Nucleic Acids Res 27:4305–13.

46. Robinson M, Son B, Kroos D, Kroos L. 2014. Transcription factor MrpC binds to promoter regions of hundreds of developmentally-regulated genes in Myxococcus xanthus. BMC genomics 15:1–23.

47. Samuels DJ, Frye JG, Porwollik S, McClelland M, Mrázek J, Hoover TR, Karls AC. 2013. Use of a promiscuous, constitutively-active bacterial enhancer-binding protein to define the σ 54 (RpoN) regulon of Salmonella Typhimurium LT2. BMC genomics 14:1–18.

48. Bono AC, Hartman CE, Solaimanpour S, Tong H, Porwollik S, McClelland M, Frye JG, Mrázek J, Karls AC. 2017. Novel DNA binding and regulatory activities for σ54 (RpoN) in Salmonella enterica serovar Typhimurium 14028s. Journal of bacteriology 199.

49. Bonocora RP, Smith C, Lapierre P, Wade JT. 2015. Genome-scale mapping of Escherichia coli σ 54 reveals widespread, conserved intragenic binding. PLoS Genet 11:e1005552.

50. Fitzgerald DM, Bonocora RP, Wade JT. 2014. Comprehensive mapping of the Escherichia coli flagellar regulatory network. PLoS Genet 10:e1004649.

51. Fitzgerald DM, Smith C, Lapierre P, Wade JT. 2018. The evolutionary impact of intragenic FliA promoters in proteobacteria. Molecular microbiology 108:361–378.

52. Mejía-Almonte C, Busby SJ, Wade JT, van Helden J, Arkin AP, Stormo GD, Eilbeck K, Palsson BO, Galagan JE, Collado-Vides J. 2020. Redefining fundamental concepts of transcription initiation in bacteria. Nature Reviews Genetics 21:699–714.

53. Gronewold TM, Kaiser D. 2002. act operon control of developmental gene expression in Myxococcus xanthus. Am Soc Microbiol.

54. Sun H, Shi W. 2001. Analyses of mrp genes during Myxococcus xanthus development. Journal of Bacteriology 183:6733–6739.

55. McLaughlin PT, Bhardwaj V, Feeley BE, Higgs PI. 2018. MrpC, a CRP/Fnr homolog, functions as a negative autoregulator during the Myxococcus xanthus multicellular developmental program. Molecular microbiology 109:245–261.

56. Wall D, Kaiser D. 1999. Type IV pili and cell motility. Molecular microbiology 32:01–10.

57. Bretl DJ, Müller S, Ladd KM, Atkinson SN, Kirby JR. 2016. Type IVupili dependent motility is couregulated by PilSR and PilS2R2 twoucomponent systems via distinct pathways in Myxococcus xanthus. Molecular microbiology 102:37–53.

58. Jinek M, Chylinski K, Fonfara I, Hauer M, Doudna JA, Charpentier E. 2012. A programmable dual-RNA–guided DNA endonuclease in adaptive bacterial immunity. science 337:816–821.

59. Heler R, Marraffini LA, Bikard D. 2014. Adapting to new threats: the generation of memory by CRISPRuCas immune systems. Molecular microbiology 93:1–9.

60. Marraffini LA. 2015. CRISPR-Cas immunity in prokaryotes. Nature 526:55–61.

61. Mojica FJ, RodriguezuValera F. 2016. The discovery of CRISPR in archaea and bacteria. The FEBS journal 283:3162–3169.

62. Jiang F, Doudna JA. 2017. CRISPR–Cas9 structures and mechanisms. Annual review of biophysics 46:505–529.

63. Shaibe E, Metzer E, Halpern YS. 1985. Control of utilization of L-arginine, L-ornithine, agmatine, and putrescine as nitrogen sources in Escherichia coli K-12. Journal of bacteriology 163:938–942.

64. Ma D, Cook DN, Alberti M, Pon NG, Nikaido H, Hearst JE. 1995. Genes acrA and acrB encode a stressuinduced efflux system of Escherichia coli. Molecular microbiology 16:45–55.

65. Bay DC, Stremick CA, Slipski CJ, Turner RJ. 2017. Secondary multidrug efflux pump mutants alter Escherichia coli biofilm growth in the presence of cationic antimicrobial compounds. Research in microbiology 168:208–221.

66. May KL, Lehman KM, Mitchell AM, Grabowicz M. 2019. A stress response monitoring lipoprotein trafficking to the outer membrane. Mbio 10:e00618–19.

67. McMahon SA, Zhu W, Graham S, Rambo R, White MF, Gloster TM. 2020. Structure and mechanism of a Type III CRISPR defence DNA nuclease activated by cyclic oligoadenylate. Nature communications 11:1–11.

68. Kim KI, Park S-C, Kang SH, Cheong G-W, Chung CH. 1999. Selective degradation of unfolded proteins by the self-compartmentalizing HtrA protease, a periplasmic heat shock protein in Escherichia coli. Journal of molecular biology 294:1363–1374.

69. Skorko-Glonek J, Zurawa D, Kuczwara E, Wozniak M, Wypych Z, Lipinska B. 1999. The Escherichia coli heat shock protease HtrA participates in defense against oxidative stress. Molecular and General Genetics MGG 262:342–350.

70. Rowley G, Spector M, Kormanec J, Roberts M. 2006. Pushing the envelope: extracytoplasmic stress responses in bacterial pathogens. Nature Reviews Microbiology 4:383–394.

71. Choi U, Lee C-R. 2019. Distinct roles of outer membrane porins in antibiotic resistance and membrane integrity in Escherichia coli. Frontiers in microbiology 10:953.

