## Supplemental Files for "Unraveling a Bacterial Starvation Response Through the Direct Targets of a Starvation-Induced Transcriptional Activator"

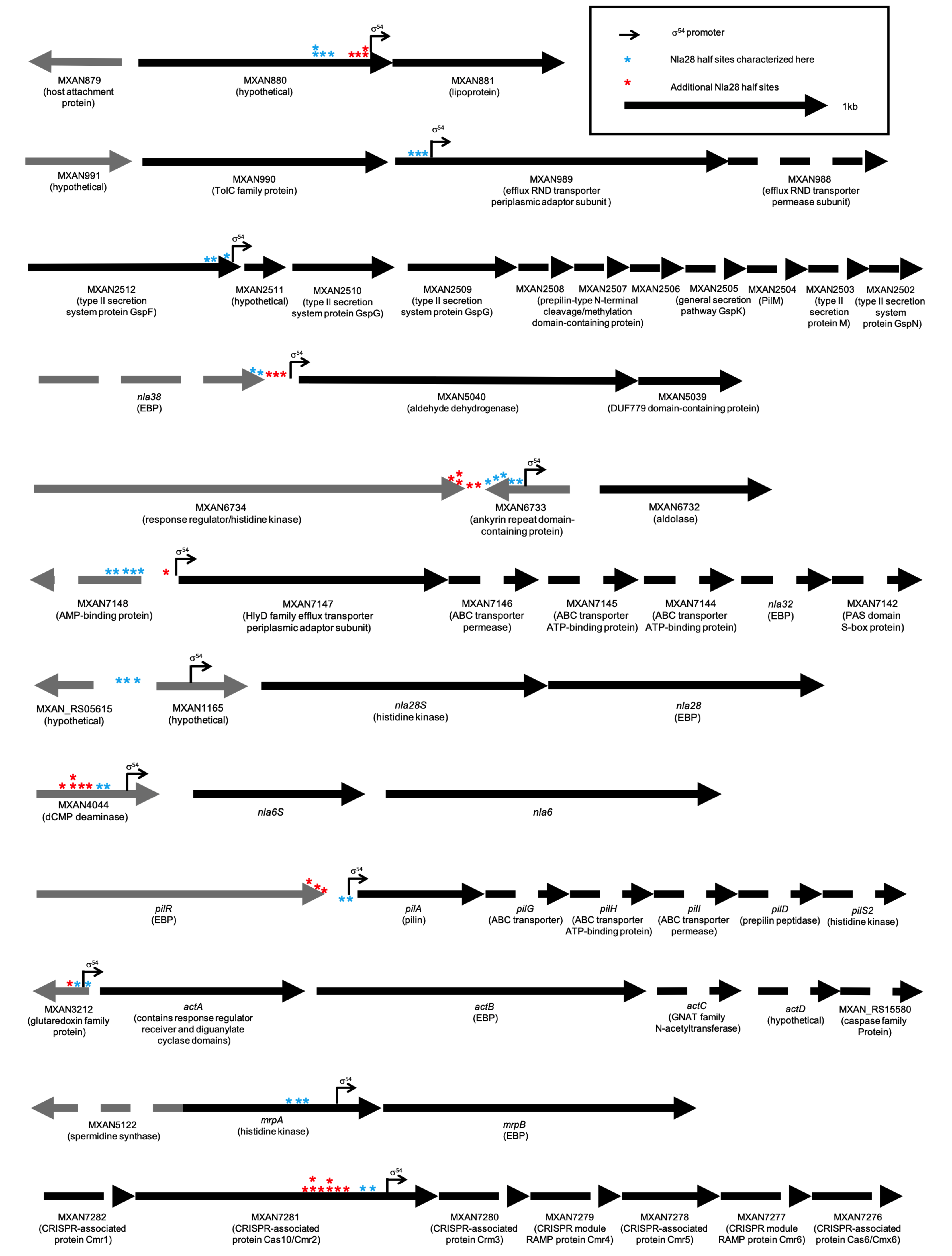

**FIGURE S1** Locations of putative Nla28 half binding sites in *actB*, *nla6*, *nla28*, *mrpB*, *pilA*, MXAN881, MXAN988, MXAN2511, MXAN5040, MXAN6732, MXAN7147, MXAN7280 promoters.

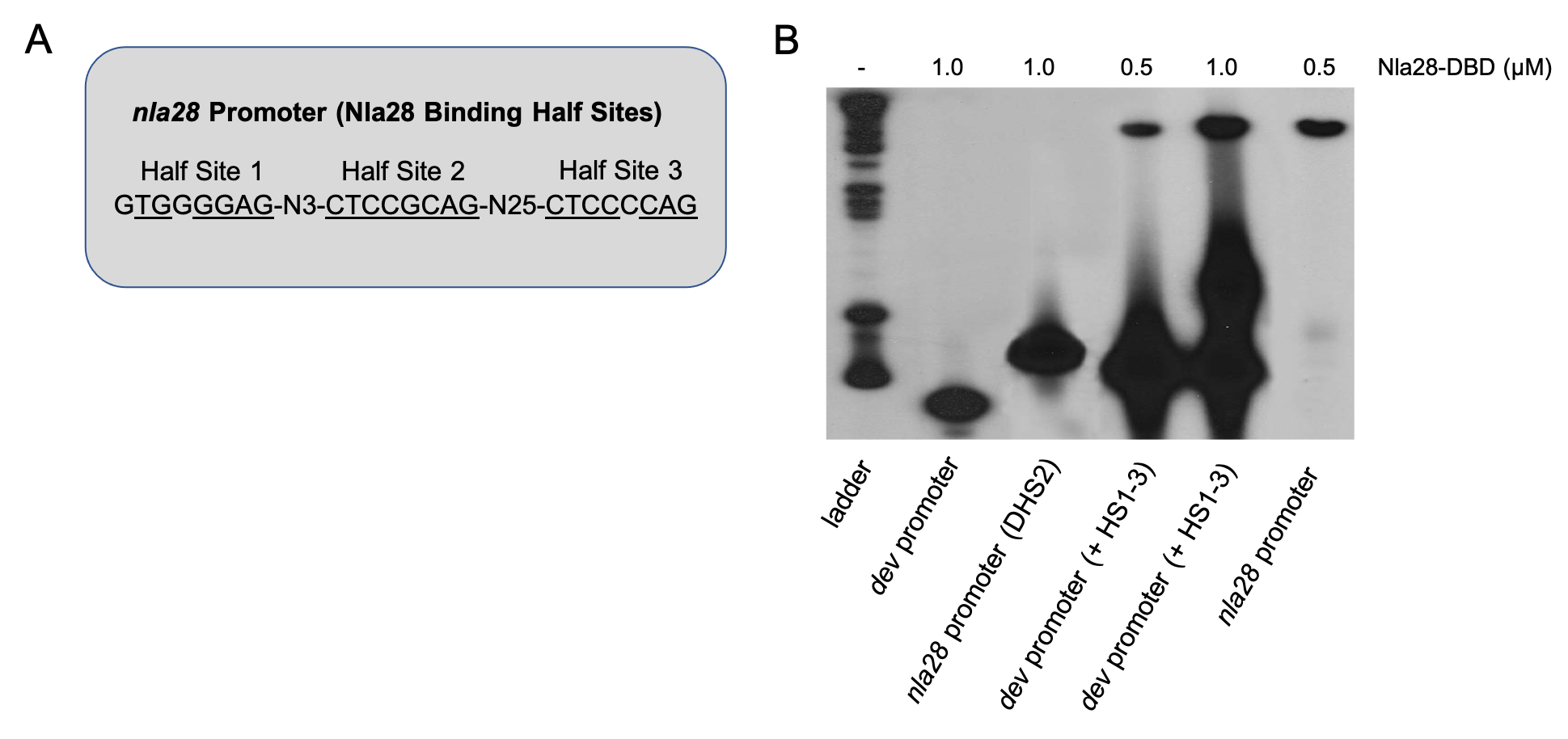

**FIGURE S2** (A) Three 8-bp sequences (Half Site 1, Half Site 2 and Half Site 3) that closely match the consensus Nla28 half binding were identified in the wild-type *nla28* promoter fragment. Underlined sequences represent nucleotides that match consensus Nla28 half binding site. (B) EMSAs were performed with 0.5 or 1.0 μM purified Nla28-DBD and 32^P^-labeled promoter fragments (wild-type dev promoter, Half Site 2 deleted *nla28* promoter (DHS2), dev promoter containing the inserted three wild-type *nla28* promoter half sites (+ HS1-3) and wild-type *nla28* promoter).

**TABLE S1** Strains and Plasmids

| Strain or Plasmid | Relevant Characteristics | Reference |
| --- | --- | --- |
| ***E***. ***coli*** **strains** |  |  |
| BL21 (DE3) | F^–^ *ompT* *gal* *dcm* *lon* *hsdS_B_*(*r_B_*^–^*m_B_*^–^) λ(DE3 [*lacI* *lacUV5*-*T7p07* *ind1* *sam7* *nin5*]) [*malB*^+^]_K-12_(λ^S^) | (1) |
| TOP10 | F- *mcrA* Δ( *mrr-hsd*RMS-*mcr*BC) Φ80*lac*ZΔM15 Δ *lac*X74 *rec*A1 *ara*D139 Δ( *araleu*)7697 *gal*U *gal*K *rps*L (StrR) *end*A1 *nup*G | Invitrogen |
| ***M*. xanthus strains** |  |  |
| DK1622 | Wild-type (parental strain) | (2) |
| AG328 | pNBC28::*nla28* (insertion in the *nla28* gene) | (3) |
| AG1251 | pTL2::*nla28* locus (*lacZ* fused to wild-type *nla28* promoter) | This study |
| AG1252 | pTL3::*nla28* locus (*lacZ* fused to mutated *nla28* promoter) | This study |
| AG1254 | pTL5::*nla28* locus (*lacZ* fused to mutated *nla28* promoter) | This study |
| AG1256 | pTL7::*nla28* locus (*lacZ* fused to mutated *nla28* promoter) | This study |
| AG1258 | pTL9::*nla28* locus (*lacZ* fused to mutated *nla28* promoter) | This study |
| AG1262 | pTL13::*attB* (*lacZ* fused to wild-type *nla28* promoter) | This study |
| AG1263 | pTL14::*attB* (*lacZ* fused to mutated *nla28* promoter) | This study |
| DK 2161 | A^-^S^-^ | (4) |
| DK10603 | D*actB* | (5) |
| MXAN881 | pKAM1001::MXAN881 (insertion in the MXAN881 gene) | This study |
| MXAN989 | pKAM1002::MXAN989 (insertion in the MXAN989 gene ) | This study |
| MXAN2510 | pKAM1003::MXAN2510 (insertion in the MXAN2510 gene) | This study |
| MXAN5040 | pKAM1004::MXAN5040 (insertion in the MXAN5040 gene) | This study |
| MXAN6732 | pEC101::MXAN6732 (insertion in the MXAN6732 gene) | This study |
| MXAN7147 | pKAM1005::MXAN7147 (insertion in the MXAN7147 gene) | This study |
| MXAN7279 | pKAM1006::MXAN7279 (insertion in the MXAN7279 gene) | This study |
| DK10410 | D*pilA* | (6) |
| SW2802 | D*mrpB* | (7) |
| **Plasmids** |  |  |
| pKAM1001 | pCR2.1TOPO (Invitrogen) containing a 319-bp fragment of the MXAN881 gene | This study |
| pKAM1002 | pCR2.1TOPO (Invitrogen) containing a 459-bp fragment of the MXAN989 gene | This study |
| pKAM1003 | pCR2.1TOPO containing a 231-bp fragment of the MXAN2510 gene | This study |
| pKAM1004 | pCR2.1TOPO containing a 364-bp fragment of the MXAN5040 gene | This study |
| pKAM1005 | pCR2.1TOPO containing a 349-bp fragment of the MXAN7147 gene | This study |
| pKAM1006 | pCR2.1TOPO containing a 377-bp fragment of the MXAN7279 gene | This study |
| pMAL-c5x | Amp^r^ (Maltose-binding protein fusion vector for Nla28-DBD protein expression) | New England Biolabs |
| pMM302 | Amp^r^ (Fragment for Nla28-DBD expression in pMAL-c5x) | (8) |
| pREG1727 | Amp^r^ Kan^r^ (attP for integration at the chromosomal Mx8 attachment site and promoterless lacZYA genes) | (9) |
| pTL1 | Amp^r^ Kan^r^ (a derivative of pREG1727 with attP removed) | This study |
| pTL2 | pTL1 with a 494-bp nla28 promoter fragment that contains the three Nla28 half binding sites (HS1-3) and the s^54^ promoter | This study |
| pTL3 | pTL2 with a 2-bp substitution in HS2 (CTCCGCAG to **AC**CCGCAG) | This study |
| pTL5 | pTL2 with a 2-bp substitution in HS2 (CTCCGCAG to CTC**AA**CAG) | This study |
| pTL7 | pTL2 with a 4-bp substitution in HS2 (CTCCGCAG to **AC**C**AA**CAG) | This study |
| pTL9 | pTL2 with a 6-bp substitution in HS2 (CTCCGCAG to **ACAAAA**AG) | This study |
| pTL13 | pREG1727 containing the 494-bp nla28 promoter fragment | This study |
| pTL14 | pTL13 with a 2-bp substitution in HS2 (CTCCGCAG to **AC**CCGCAG) | This study |
| pTL27 | pCR-Blunt (Invitrogen) containing the 494-bp nla28 promoter fragment | This study |
| pTL28 | pTL27 with a 2-bp substitution in HS2 (CTCCGCAG to **AC**CCGCAG) | This study |
| pTL30 | pTL27 with a 2-bp substitution in HS2 (CTCCGCAG to CTC**AA**CAG) | This study |
| pTL32 | pTL27 with a 4-bp substitution in HS2 (CTCCGCAG to **AC**C**AA**CAG) | This study |
| pTL34 | pTL27 with a 6-bp substitution in HS2 (CTCCGCAG to **ACAAAA**AG) | This study |

References

1.	Studier FW, Moffatt BA. 1986. Use of bacteriophage T7 RNA polymerase to direct selective high-level expression of cloned genes. Journal of molecular biology 189:113-130.

2.	Kaiser D. 1979. Social gliding is correlated with the presence of pili in Myxococcus xanthus. Proceedings of the National Academy of Sciences 76:5952-5956.

3.	Caberoy NB, Welch RD, Jakobsen JS, Slater SC, Garza AG. 2003. Global mutational analysis of NtrC-like activators in Myxococcus xanthus: identifying activator mutants defective for motility and fruiting body development. Journal of Bacteriology 185:6083-6094.

4.	Hodgkin J, Kaiser D. 1979. Genetics of gliding motility in Myxococcus xanthus (Myxobacterales): two gene systems control movement. Molecular and General Genetics MGG 171:177-191.

5.	Gronewold TM, Kaiser D. 2001. The act operon controls the level and time of C‐signal production for Myxococcus xanthus development. Molecular microbiology 40:744-756.

6.	Wu SS, Kaiser D. 1996. Markerless deletions of pil genes in Myxococcus xanthus generated by counterselection with the Bacillus subtilis sacB gene. Journal of Bacteriology 178:5817-5821.

7.	Sun H, Shi W. 2001. Genetic studies of mrp, a locus essential for cellular aggregation and sporulation of Myxococcus xanthus. Journal of bacteriology 183:4786-4795.

8.	Ma M, Welch RD, Garza AG. 2021. The σ 54 system directly regulates bacterial natural product genes. Scientific reports 11:1-11.

9.	Fisseha M, Gloudemans M, Gill RE, Kroos L. 1996. Characterization of the regulatory region of a cell interaction-dependent gene in Myxococcus xanthus. Journal of Bacteriology 178:2539-2550.

**TABLE S2** Primers

|  | Sequence | Amplicon size |
| --- | --- | --- |
| **qPCR** |  |  |
| *actB* foward  *actB* reverse | 5’-CTCCAGGACGAGGAGTTCTTCCG-3’  5’-GCGATTTCCTTCTCCAGGTCGC-3’ | 101 bp |
| *mrpB* forward  *mrpB* reverse | 5’-GCCAGCCTGATTCCCACTTT-3’  5’-ACCGTACTTCTGGAGCTTGC-3’ | 144 bp |
| *nla6* forward  *nla6* reverse | 5’-GCTCATCGAGTCCGAGCTG-3’  5’-ATTCGTCCAGGAAGAGCGTG-3’ | 113 bp |
| *nla28* forward  *nla28* reverse | 5’-GTGTTGCAGGAGGGCGAAATCC-3’  5’-CGCGTCTTGAGGTCCTTGTTCG-3’ | 98 bp |
| *nla28S* (MXAN1166) forward  *nla28S* (MXAN1166) reverse | 5'-CGCTCGGAAGAAGGAAAGGG-3'  5'-CGGAAGCAGTCGTTGTCTGA-3' | 100 bp |
| *pilA* forward  *pilA* reverse | 5'-GTCGTTGGAGATGCACGGAACG-3'  5'-CAACCGTTACGGCTACCGTGTG-3' | 106 bp |
| *rpoD* forward  *rpoD* reverse | 5’-GACGTCTTGGAGCGAGAGCTGTC-3’  5’-CTCCATCATCTTGGTCCGGAGGTC-3’ | 102 bp |
| MXAN881foward  MXAN881reverse | 5’-ATGATGGGGATGGCTTGGTG-3’  5’-GCAGCGCGGATAATCGTTTT-3’ | 104 bp |
| MXAN989 forward  MXAN989 reverse | 5’-GGAGAAGAACGTGGTGGAGG-3’  5’-GGTCCTGCTCGTACACATCC-3’ | 107 bp |
| MXAN2511 forward  MXAN2511 reverse | 5’-TGGCGGAATGATGAACGACA-3’  5’-TTGCTGGGGACCGTACAAC-3’ | 118 bp |
| MXAN5040 forward  MXAN5040 reverse | 5’-GTGGACCAACTGCTACCACA-3’  5’-TGACCAGCAGGTTCTTCGTC-3’ | 128 bp |
| MXAN6732 forward  MXAN6732 reverse | 5’-ATGACTGCGCCTTCCTGAAA-3’  5’-GTTGACCGTGATGGGAGAGG-3’ | 117 bp |
| MXAN7147 forward  MXAN7147 reverse | 5’-TTCACCGTGTCCAACCACAC-3’  5’-TGGACATTCCCAAAGCCAAGA-3’ | 154 bp |
| MXAN7279 forward  MXAN7279 reverse | 5'-GCCACGGGTACCTTCAACAA-3'  5'-GGCCTTGCCTCCCAGTTG-3' | 105 bp |
| **Promoter fragments** |  |  |
| *actB* forward  *actB* reverse | 5’-CCGCCGTCGTGGAGTC-3’  5’-CGGATTTAGCAATGGTTGTGCCAC-3’ | 210 bp |
| *dev* forward  *dev* reverse | 5’-ACGTTGCAGACGGGGTGAG-3’  5’-CCTCGTACTTCGACTTCCGAAGAG-3’ | 180 bp |
| *mrpB* forward  *mrpB* reverse | 5’-TGAGGCCGGTGTTCCGGG-3’  5’-TCCGCCAACCTCGGCGG-3’ | 220 bp |
| *nla28* forward  *nla28* reverse | 5’-GATGACGCGCGCAGCTTGC-3’  5’-GAGATTGCGCCGTCCCAGC-3’ | 200 bp |
| *pilA* forward  *pilA* reverse | 5’-GAGCGCTTCGGATGCGTAGG-3’  5’-TCCTCAGAGAAGGTTGCAACGG-3’ | 200 bp |
| MXAN881 forward  MXAN881 reverse | 5’-CGCGCTTCTCCACGTCCTTG-3’  5’-GACGTCATGCTGGAGATTTCCGG-3’ | 196 bp |
| MXAN989 forward  MXAN989 reverse | 5’-GTAATCGTCGCCGTGGCCTG-3’  5’-TCCTTCCAGGGTGTCCTGCG-3’ | 201 bp |
| MXAN2511 forward  MXAN2511 reverse | 5’-CTCCGCTGGTGTACCACATGG-3’  5’-CAGAATCGGCATCAGGATGGAGA-3’ | 200 bp |
| MXAN5040 forward  MXAN5040 reverse | 5’-GTCCGTTGCGCGAGCTGGA-3’  5’-CGCGACAGGTGCGTCGC-3’ | 200 bp |
| MXAN6732 forward  MXAN6732 reverse | 5’-GGGGCTGTCTCCGTGACGC-3’  5’-CGCAACGGGCATGCCGA-3’ | 201 bp |
| MXAN7147 forward  MXAN7147 reverse | 5’-AGGTGATGGCGAAGCGGGC-3’  5’-CCTTCTCGAAGTGTTCCTGGACCA-3’ | 197 bp |
| MXAN7280 forward  MXAN7280 reverse | 5’-CAGTGCTGGTGGCGAAGCAC-3’  5’-CTCAGCCAGCTCGGGCG-3’ | 200 bp |
| **Insertions** |  |  |
| MXAN881 forward  MXAN881 reverse | 5’-CTGCACGTCCCTGCCGGAC-3’  5’-GTTGCAGGCGCCGCAGTGG-3’ | 319 bp |
| MXAN989 forward  MXAN989 reverse | 5’-CTGCTGGCGCGTCAGAGC-3’  5’-GCGGACGTCCGCGTACATG-3’ | 459 bp |
| MXAN2510 forward  MXAN2510 reverse | 5’-GAAGCAGCGCCGCAACCG-3’  5’-GCCTCCACCAGCGCTTGC-3’ | 231 bp |
| MXAN5040 forward  MXAN5040 reverse | 5’-GAGCAGACGCCCTCCAGC-3’  5’-GCTCGCTGATGAGGGAGCGC-3’ | 364 bp |
| MXAN6732 foward  MXAN6732 reverse | 5’-GGATGGCCAACCCGGCGA-3’  5’-AAGATCCAGTCGTGCGCCTCC-3’ | 418 bp |
| MXAN7147 forward  MXAN7147 reverse | 5’-GCCGAGCGTCAACTGGCC-3’  5’-TCGACTCCACCTGCGCGC-3’ | 349 bp |
| MXAN7279 forward  MXAN7279 reverse | 5’-CGAAGGCGGGTCCGGAGG-3’  5’-CTCCGCCTCCGGCAGCAG-3’ | 377 bp |

**TABLE S4** Potential targets of Nla28 in *M. xanthus*

| Locus | Number of genes | Peak expression  time point  in WT cells (h) | Fold increase of expression at peak time point in WT cells | Fold decrease of expression  in *nla28* ^-^ cells |
| --- | --- | --- | --- | --- |
| MXAN162 | 1 | 12 | 2.5 | 8.3 |
| MXAN179 | 3 | 24 | 6.1 | 3.7 |
| MXAN255 | 1 | 12 | 2.2 | 4.2 |
| MXAN419 | 2 | 12 | 2.1 | 5.6 |
| MXAN496 | 2 | 12 | 3.1 | 3.7 |
| MXAN542 | 2 | 12 | 2.3 | 3.7 |
| MXAN562 | 1 | 1 | 2.3 | 3.6 |
| MXAN854 | 1 | 6 | 4.1 | 3.0 |
| MXAN859 | 1 | 12 | 3.1 | 2.5 |
| MXAN909 | 2 | 12 | 2.4 | 3.3 |
| MXAN934 | 1 | 6 | 2.6 | 3.7 |
| MXAN1043 | 2 | 12 | 2.1 | 6.7 |
| MXAN1124 | 1 | 12 | 2.4 | 2.4 |
| MXAN1404 | 1 | 12 | 2.2 | 4.3 |
| MXAN1412 | 1 | 6 | 3.1 | 3.6 |
| MXAN1433 | 1 | 1 | 2.1 | 3.3 |
| MXAN1752 | 2 | 6 | 2.7 | 3.3 |
| MXAN1783 | 1 | 12 | 2.0 | 2.9 |
| MXAN1968 | 1 | 1 | 3.2 | 1.7 |
| MXAN2159 (*nla12*) | 2 | 12 | 2.4 | 4.3 |
| MXAN2359 | 2 | 24 | 2.4 | 4.8 |
| MXAN2362 | 1 | 12 | 2.8 | 2.0 |
| MXAN2363 | 1 | 12 | 2.8 | 2.4 |
| MXAN2447 | 7 | 24 | 2.4 | 2.9 |
| MXAN2515 | 5 | 24 | 3.1 | 2.9 |
| MXAN2735 | 1 | 12 | 2.0 | 5.3 |
| MXAN2752 | 2 | 12 | 2.3 | 2.6 |
| MXAN2777 | 1 | 12 | 2.2 | 7.1 |
| MXAN2907 | 1 | 12 | 2.3 | 2.6 |
| MXAN2947 | 1 | 12 | 2.7 | 7.7 |
| MXAN2952 | 5 | 6 | 2.2 | 1.8 |
| MXAN3026 | 2 | 6 | 3.7 | 5.9 |
| MXAN3149 | 2 | 6 | 9.0 | 5.3 |
| MXAN3167 | 1 | 12 | 2.1 | 3.8 |
| MXAN3735 | 2 | 12 | 2.1 | 2.2 |
| MXAN3778 | 1 | 12 | 2.2 | 4.3 |
| MXAN3824 | 1 | 12 | 2.6 | 2.7 |
| MXAN3962 | 2 | 12 | 2.8 | 2.3 |
| MXAN3975 | 1 | 12 | 2.6 | 3.1 |
| MXAN3979 | 2 | 12 | 2.3 | 8.3 |
| MXAN4007 | 1 | 1 | 2.9 | 2.3 |
| MXAN4155 | 1 | 12 | 2.2 | 5.9 |
| MXAN4198 | 4 | 18 | 2.0 | 3.8 |
| MXAN4236 | 1 | 6 | 2.3 | 1.7 |
| MXAN4687 | 1 | 1 | 7.5 | 4.5 |
| MXAN4784 | 2 | 6 | 2.0 | 2.8 |
| MXAN4877 | 1 | 2 | 2.6 | 2.1 |
| MXAN5543 | 6 | 6 | 4.3 | 2.3 |
| MXAN5615 | 1 | 6 | 2.1 | 5.0 |
| MXAN5715 | 1 | 6 | 3.3 | 100.0 |
| MXAN6209  (*sigC*) | 1 | 1 | 6.1 | 5.3 |
| MXAN6223 | 5 | 6 | 4.3 | 2.1 |
| MXAN6297 | 1 | 12 | 2.5 | 4.0 |
| MXAN6437 | 1 | 6 | 2.6 | 2.2 |
| MXAN6788 | 1 | 6 | 3.2 | 33.3 |
| MXAN7157 | 1 | 18 | 2.0 | 2.9 |
| MXAN7225 | 2 | 12 | 2.4 | 5.0 |
| MXAN7435 | 1 | 1 | 4.6 | 2.0 |

**TABLE S5** Functional categories of confirmed Nla28 targets in *M. xanthus*

| Locus | Number of genes | NCBI gene definition | NCBI Cluster of Orthologous Genes (COG) Category: |
| --- | --- | --- | --- |
| *actB* | 4 | sigma-54-dependent Fis family transcriptional regulator | Signal transduction mechanisms |
| *nla6* | 2 | NtrC family HydH-HydG (metal tolerance) | Signal transduction mechanisms |
| *nla28* | 2 | NtrC family AtoS-AtoC (cPHB biosynthesis) | Signal transduction mechanisms |
| *mrpB*  (MXAN5124) | 1 | sigma-54 dependent DNA-binding response regulator | Signal transduction mechanisms |
| *pilA*  (MXAN5783) | 6 | Bacterial motility proteins | Cell motility/ Extracellular Structures |
| MXAN881 | 1 | Hypothetical lipoprotein | Poorly Characterized |
| MXAN989 | 2 | cation efflux system protein CusB | Inorganic ion transport and metabolism |
| MXAN2511 | 10 | hypothetical protein | Poorly Characterized |
| MXAN5040 | 2 | aldehyde dehydrogenase family protein | Lipid transport and metabolism |
| MXAN6732 | 1 | class II aldolase/adducin domain protein | Carbohydrate transport and metabolism/ Amino acid transport and metabolism |
| MXAN7147 | 2 | efflux transporter, RND family, MFP subunit | Defense mechanisms |
| MXAN7280 | 5 | putative CRISPR-associated protein Crm3 | Defense mechanisms |

**TABLE S6** Functional categories of other potential Nla28 targets in *M. xanthus*

| Locus | Number of genes | NCBI definition | NCBI Cluster of Orthologous Genes (COG) Category: |
| --- | --- | --- | --- |
| MXAN162 | 1 | hypothetical protein | Poorly Characterized |
| MXAN179 | 3 | metallo-beta-lactamase family protein | Poorly Characterized |
| MXAN255 | 1 | peptidase homolog, M20 family | Amino acid transport and metabolism |
| MXAN419 | 2 | hypothetical protein | Signal transduction mechanisms |
| MXAN496 | 2 | glyoxalase family protein | General function prediction only |
| MXAN542 | 2 | F5/8 type C domain protein | Poorly Characterized |
| MXAN562 | 1 | phosphate-selective porin O and P | Poorly Characterized |
| MXAN854 | 1 | hypothetical protein. | Poorly Characterized |
| MXAN859 | 1 | putative lipoprotein | Poorly Characterized |
| MXAN909 | 2 | hypothetical protein | Poorly Characterized |
| MXAN934 | 1 | protease DO family protein | Posttranslational modification, protein turnover, chaperones |
| MXAN1043 | 2 | glycosyl transferase | Cell wall/membrane/envelope biogenesis |
| MXAN1124 | 1 | ABC transporter, ATP-binding protein | Cell wall/membrane/envelope biogenesis |
| MXAN1404 | 1 | hypothetical protein TIGR00661 | Poorly Characterized |
| MXAN1412 | 1 | putative serine/threonine protein phosphatase | Signal transduction mechanisms |
| MXAN1433 | 1 | M23 peptidase domain protein | Cell wall/membrane/envelope biogenesis |
| MXAN1752 | 2 | isoquinoline 1-oxidoreductase, beta subunit | Energy production and conversion |
| MXAN1783 | 1 | hypothetical protein | Poorly Characterized |
| MXAN1968 | 1 | hypothetical protein | Poorly Characterized |
| MXAN2159 (*nla12*) | 2 | sigma-54 dependent transcriptional regulator, Fis family | Signal transduction mechanisms |
| MXAN2359 | 2 | glycosyl transferase, group 2 family protein | Cell wall/membrane/envelope biogenesis |
| MXAN2362 | 1 | glycosyl transferase, group 1 family protein | Cell wall/membrane/envelope biogenesis |
| MXAN2363 | 1 | glutaminyl-tRNA synthetase | Translation, ribosomal structure and biogenesis |
| MXAN2447 | 7 | FliP/MopC/SpaP family protein | Intracellular trafficking, secretion, and vesicular transport |
| MXAN2515 | 5 | general secretion pathway protein C | Intracellular trafficking, secretion, and vesicular transport |
| MXAN2735 | 1 | methyl-accepting chemotaxis protein | Cell motility |
| MXAN2752 | 2 | hypothetical protein | Poorly Characterized |
| MXAN2777 | 1 | tonB domain protein | Cell wall/membrane/envelope biogenesis |
| MXAN2907 | 1 | CDP-diacylglycerol--serine O-phosphatidyltransferase | Lipid transport and metabolism |
| MXAN2947 | 1 | isochorismatase family protein | Coenzyme transport and metabolism |
| MXAN2952 | 5 | hypothetical protein | Poorly Characterized |
| MXAN3026 | 2 | O-antigen polymerase family protein | Cell wall/membrane/envelope biogenesis |
| MXAN3149 | 2 | peptidase, M6 (immune inhibitor A) family | Posttranslational modification, protein turnover, chaperones |
| MXAN3167 | 1 | hypothetical protein | Poorly Characterized |
| MXAN3735 | 2 | response regulator | Transcription/Signal transduction mechanisms |
| MXAN3778 | 1 | DnaK family protein | Posttranslational modification, protein turnover, chaperones |
| MXAN3824 | 1 | general secretion pathway protein G | Cell motility/ Extracellular structures/ Intracellular trafficking, secretion, and vesicular transport |
| MXAN3962 | 2 | pyridine nucleotide-disulphide oxidoreductase | Amino acid transport and metabolism |
| MXAN3975 | 1 | hypothetical protein | Poorly Characterized |
| MXAN3979 | 2 | hypothetical protein | Poorly Characterized |
| MXAN4007 | 1 | hypothetical protein | Poorly Characterized |
| MXAN4155 | 1 | hypothetical protein | Poorly Characterized |
| MXAN4198 | 4 | putative outer membrane macrolide efflux protein | Cell wall/membrane/envelope biogenesis |
| MXAN4236 | 1 | CBS domain protein | Signal transduction mechanisms |
| MXAN4687 | 1 | YHS domain protein | Inorganic ion transport and metabolism |
| MXAN4784 | 2 | inorganic anion transporter, sulfate permease (SulP) family | Inorganic ion transport and metabolism |
| MXAN4877 | 1 | aspartate kinase | Amino acid transport and metabolism |
| MXAN5543 | 6 | ATPase, P-type (transporting), HAD superfamily, subfamily IC | Inorganic ion transport and metabolism |
| MXAN5615 | 1 | hypothetical protein | Poorly Characterized |
| MXAN5715 | 1 | response regulator/putative sensor histidine kinase | Signal transduction mechanisms |
| MXAN6209 (*sigC*) | 1 | RNA polymerase sigma-C factor | Transcription |
| MXAN6223 | 5 | sensor histidine kinase | Signal transduction mechanisms |
| MXAN6297 | 1 | ribulose-phosphate 3-epimerase | Carbohydrate transport and metabolism |
| MXAN6437 | 1 | putative lipoprotein | Poorly Characterized |
| MXAN6788 | 1 | conserved domain protein | Poorly Characterized |
| MXAN7157 | 1 | enoyl-CoA hydratase/isomerase family protein | Lipid transport and metabolism |
| MXAN7225 | 2 | putative sugar ABC transporter, ATP-binding protein | Carbohydrate transport and metabolism |
| MXAN7435 | 1 | hydrolase, alpha/beta fold family | Coenzyme transport and metabolism |
